# Feeding-mode-defined microbial communities modulate prebiotic responses and alters colonic motility in early life

**DOI:** 10.64898/2026.05.29.728681

**Authors:** Markus Schwalbe, Arjen Nauta, Ellen Looijesteijn, Fatima Pereira, Sahar El Aidy

## Abstract

Prebiotic supplementation a well-established impact on early-life gut microbiota, yet how substrate type and baseline community structure interact to shape microbial and metabolic responses remains incompletely understood. Using donor-derived infant faecal fermentations, a defined synthetic microbial community (SynCom), and multi-layered metabolomics, we demonstrate that prebiotic responses are strongly dependent on feeding-mode and mediated by species-specific metabolic interactions. While both human milk oligosaccharides (HMOs) and galacto-oligosaccharides (GOS) robustly promoted bifidobacterial expansion, modulation of *Bacteroides/Phocaeicola* species varied according to feeding background and substrate identity with 2′-fucosyllactose (2′FL) selectively alleviating growth inhibition in breast milk-derived communities. Mechanistic dissection using SynCom revealed extensive cross-feeding networks and substrate-dependent interaction rewiring, identifying *Phocaeicola vulgatus* as a context-dependent ecological hub that disproportionately shaped succinate and propionate production and altered *Escherichia coli* metabolic activity without affecting its abundance. Competitive outcomes were determined not only by environmental acidification, but also by the chemical identity of fermentation products, with lactate exerting species-specific inhibitory effects independent of bulk pH changes. Using an in-house *ex vivo* IntestiFlow gut organ platform, we further show that metabolites produced by the defined community induced a strong but reversible suppression of colonic peristaltic activity, an effect not recapitulated by SCFAs alone. Biogenic amine metabolites, including phenethylamine, partially reproduced key aspects of this response, suggesting a role for specific microbial fermentation products in modulating intestinal motility. Together, these findings demonstrate that early-life prebiotic responses extend beyond uniform bifidogenic expansion and instead emerge from interaction-driven metabolic specialization within feeding-mode-specific community contexts, with direct functional consequences for infant intestinal physiology.

## Introduction

The infant gut microbiome plays a central role in early-life development, influencing metabolism, immunity, and long-term health outcomes. Disruptions in early colonization, caused by delivery mode, antibiotic exposure, or household environment, have been linked to increased risks of allergies, asthma, and obesity during childhood (Azad et al., 2014; Darabi et al., 2019; Hoskinson et al., 2023). Infants delivered by C-section, for example, exhibit reduced colonisation by *Bacteroides* and *Bifidobacterium* strains and increased prevalence of potentially pathogenic *Enterococcus*, *Enterobacter* and *Klebsiella* species during the first months of life (Shao et al., 2019). Feeding mode is another major determinant of early gut microbial composition (Casaburi et al., 2021; Ho et al., 2018). Human milk oligosaccharides (HMOs), abundant in breast milk, serve as primary energy source for early colonizers, particularly *Bifidobacterium* species (Milani, Duranti, Bottacini, Casey, Turroni, Mahony, Belzer, Delgado Palacio, Arboleya Montes, Mancabelli, Lugli, Rodriguez, Bode, de Vos, et al., 2017). HMO composition can vary greatly between individuals due to genetic, e.g. secreter status, and environmental factors, with 2’-fucosylactose (2’FL) being the most prevalent HMO, spanning concentrations from 1.65 to 3.18 g/L in breast milk (Soyyılmaz et al., 2021). In contrast, formula-fed infants harbour lower abundance of Bifidobacteria, higher alpha diversity, and a microbial composition more reminiscent of adults (Casaburi et al., 2021; Ho et al., 2018), along with reduced representation of pathways related to lipid metabolism, detoxification, and vitamin synthesis. These compositional changes frequently alter production of microbial metabolites, including short chain fatty acids (SCFAs) and amino acid-derived compounds, which can impact energy metabolism and neurodevelopment (Needham et al., 2022; Tsukuda et al., 2021). Yet, most studies focus on compositional shifts, leaving unresolved how prebiotics alter metabolic interactions, cross-feeding pathways, and the downstream bioactive metabolites that shape early-life physiology.

Only a subset of gut bacteria can metabolize the full spectrum of HMOs though the infant gut hosts dozens of microbial species (Milani, Duranti, Bottacini, Casey, Turroni, Mahony, Belzer, Delgado Palacio, Arboleya Montes, Mancabelli, Lugli, Rodriguez, Bode, De Vos, et al., 2017), suggesting that cross-feeding and metabolic cooperation are critical for community functioning. Perturbations, such as transitions between breast and formula feeding may therefore disrupt these networks, with consequences for community structure, metabolite output, and host physiology. Addressing these aspects require moving beyond compositional profiling to interrogate how feeding mode reorganizes (sub)-species interactions and their metabolite outputs.

To fill these knowledge gaps, we combined *in vitro* fermentation of donor-derived infant faecal samples, with metagenomic and metabolomic profiling and a defined infant-gut microbial community, as well as an *ex vivo* gut organ system (IntestiFlow platform; **Supplementary text, Supplementary Figure S0)**, to investigate how prebiotic supplementation rewires microbial interactions, alters metabolic activity, modulates community-level metabolite outputs in a feeding-mode-dependent manner, and potentially impact infant gut motility.

## Results

### Feeding mode establishes distinct ecological baselines in infant gut communities that shape prebiotic responses

To investigate how prebiotics commonly added to infant formula (galactooligosaccharides (GOS), 2’fucosyllactose (2’FL), lacto-N-tetraose (LNT) and a 4:1 GOS:2’FL mixture) influence early-life gut communities, faecal samples from five exclusively breast-fed (BF) and five exclusively formula-fed (FF) infants were cultured individually in mini-bioreactors using SIFR technology^®^ for up to 24 h (**Figure 1A**) (Van Den Abbeele et al., 2023). Fermentation parameters (gas production, pH), metagenomic sequencing, and metabolomics were employed to quantify microbial community composition and metabolic outcomes.

**Figure 1.**
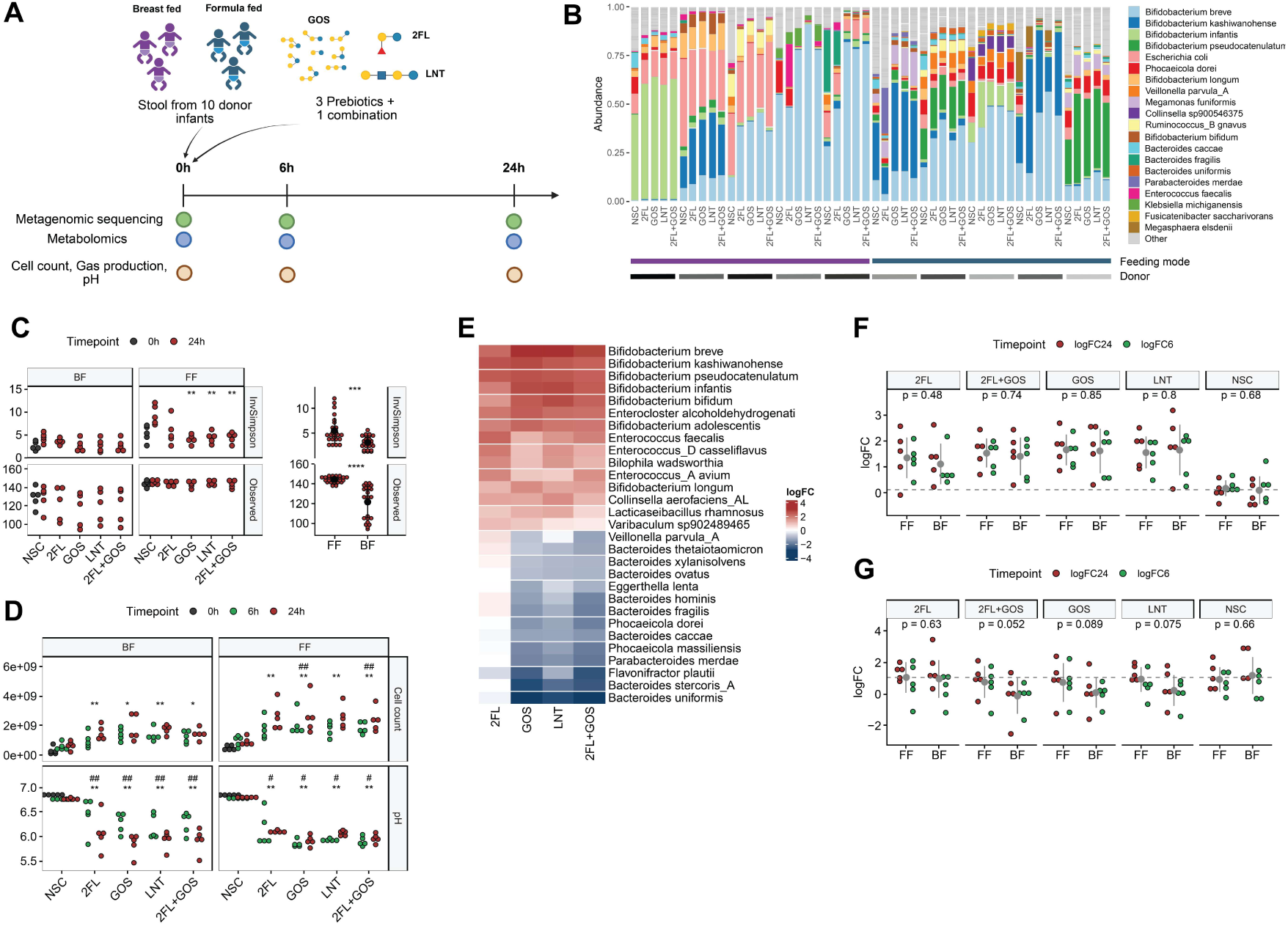
Prebiotic supplementation alters infant gut communities with feeding-mode-dependent effects on Bacteroides. (**A**) Study design. Faecal samples from ten donor infants (five breast- fed (BF) and five formula-fed (FF)) were cultured in vitro in SIFR reactors with (the combination of) galactooligosacharides (GOS), 2-fucosyllactose (2’FL) or Lacto-*N*-tetraose (LNT). Samples for metagenomics, metabolomics and other fermentation parameters were collected at 0h, 6h and 24h. (**B**) Relative microbial composition across donor, grouped by prebiotic treatment and feeding mode (top 20 species shown). (**C**) Alpha diversity (inverse Simpson index; observed number of species) showing lower diversity and strong Bifidobacterium dominance in BF communities, and higher diversity and evenness in FF communities. Prebiotic supplementation reduced evenness in FF for GOS and LNT, but not 2’FL and had a minimal impact in BF. (**D**) Fermentation parameters showing increased metabolic activity under all prebiotic conditions. (**E**) Log-fold changes in species abundance normalised to non-substrate control (NSC), highlighting consistent bifidogenic responses and selective suppression of *Bacteroides/Phocaeicola* species across most prebiotics. (**F, G**) Temporal growth dynamics of *Bifidobacterium* genus **(F)**, and *Bacteroides and Phocaeicola* genus **(G)** expressed as log-fold change relative to 0h, showing differential responses depending on feeding mode and prebiotic type.

Across all samples, 151 species (95% genome identity) were detected; at baseline (0h), communities were dominated by *Bifidobacterium* and *Bacteroides* species, with *Escherichia coli* being more prevalent in BF infants (**Figure 1B, Supplementary Figure S1A**). Although inter-individual variability was substantial, the 20 most abundant taxa accounted for about 75% in each sample. BF communities exhibited lower alpha diversity (both observed species and inverse Simpson index) and were frequently dominated by a single *Bifidobacterium* species (**Figure 1B**), whereas FF communities harboured higher diversity and evenness. Overall, feeding mode defined distinct baseline ecological configurations prior to prebiotic supplementation.

Diversity was minimally altered in BF microbial community, but evenness decreased in FF microbial community (**Figure 1C**), indicating that the microbial community with higher initial diversity underwent stronger selective expansion under prebiotic supplementation. Prebiotic supplementation increased total cell counts and gas production, while acidifying pH across both feeding modes (**Figure 1D, Supplementary Figure S1B**), consistent with enhanced microbial growth and fermentation activity.

To account for compositional effects, relative sequencing data were normalised using absolute cell counts. Across feeding modes, all prebiotic supplementations consistently stimulated growth of *Bifidobacterium* and *Enterococcus* species (**Figure 1E, Supplementary Figure S1C**). In contrast, abundance of *Bacteroides* and *Phocaeicola* species decreased under GOS, LNT, and GOS+2’FL conditions, while 2’FL selectively alleviated this suppression.

When compared to the initial inoculum, *Bifidobacterium* expansion occurred across feeding modes under all prebiotic conditions (**Figure 1F**). Conversely, the growth of *Bacteroides* and *Phocaeicola* strongly declined in BF community supplemented with LNT, GOS, or GOS+2’FL (**Figure 1G**), while 2’FL supplementation partially restored their abundance. Genomic analysis revealed that *Bacteroides* and *Phocaeicola* species encode GH29 and GH95 fucosidases (**Supplementary Excel Sheet**), suggesting that enzymatic capacity might underly substrate persistence. Taken together, these findings indicate that feeding mode establishes distinct baseline gut community structures that modulate the extent of prebiotic-driven microbiota restructuring. Although all prebiotics consistently promoted bifidobacterial expansion, 2’FL additionally supported the persistence of specific *Bacteroides/Phocaeicola* populations across different baseline community configurations. These baseline (seeding) differences appear to influence the magnitude, rather than the direction of substrate-specific microbial responses.

### Prebiotics supplementation redirects metabolic profiles and enhances propionate production in a substrate-specific manner

Analysis of short-chain fatty acids (SCFAs) revealed clear feeding-mode differences in fermentation dynamics. BF microbial communities exhibited slower SCFA accumulation compared to FF communities, consistent with lower total cell count and reduced pH at 6 h (**Figure 2A, Figure 1D**). BF communities predominantly accumulated acetate and lactate at both 6h and 24h. In contrast, FF communities showed rapid lactate consumption after 6h, followed by increased butyrate, propionate, valerate, and higher BCFAs by 24h. These differences were also visible at 0h in the initial inoculum (**Supplementary Figure S2A**). These patterns indicate that feeding mode establishes distinct baseline fermentation trajectories and lactate-utilizing capacities, consistent with its role in shaping infant gut community structure. Across donors, prebiotic supplementation increased total SCFA concentrations and reduced BCFAs levels relative to non-substrate controls (NSC), particularly in FF communities, consistent with a shift away from amino acid fermentation toward carbohydrates-driven metabolism **(Figure 2A)**. Notably, 2’FL supplementation enhanced propionate production at 24h in both feeding modes, reaching statistical significance in FF community (BF: p=0.19; FF: p=0.018). This effect was not observed with GOS or LNT, indicating substrate-specific engagement of propionate-producing populations.

**Figure 2.**
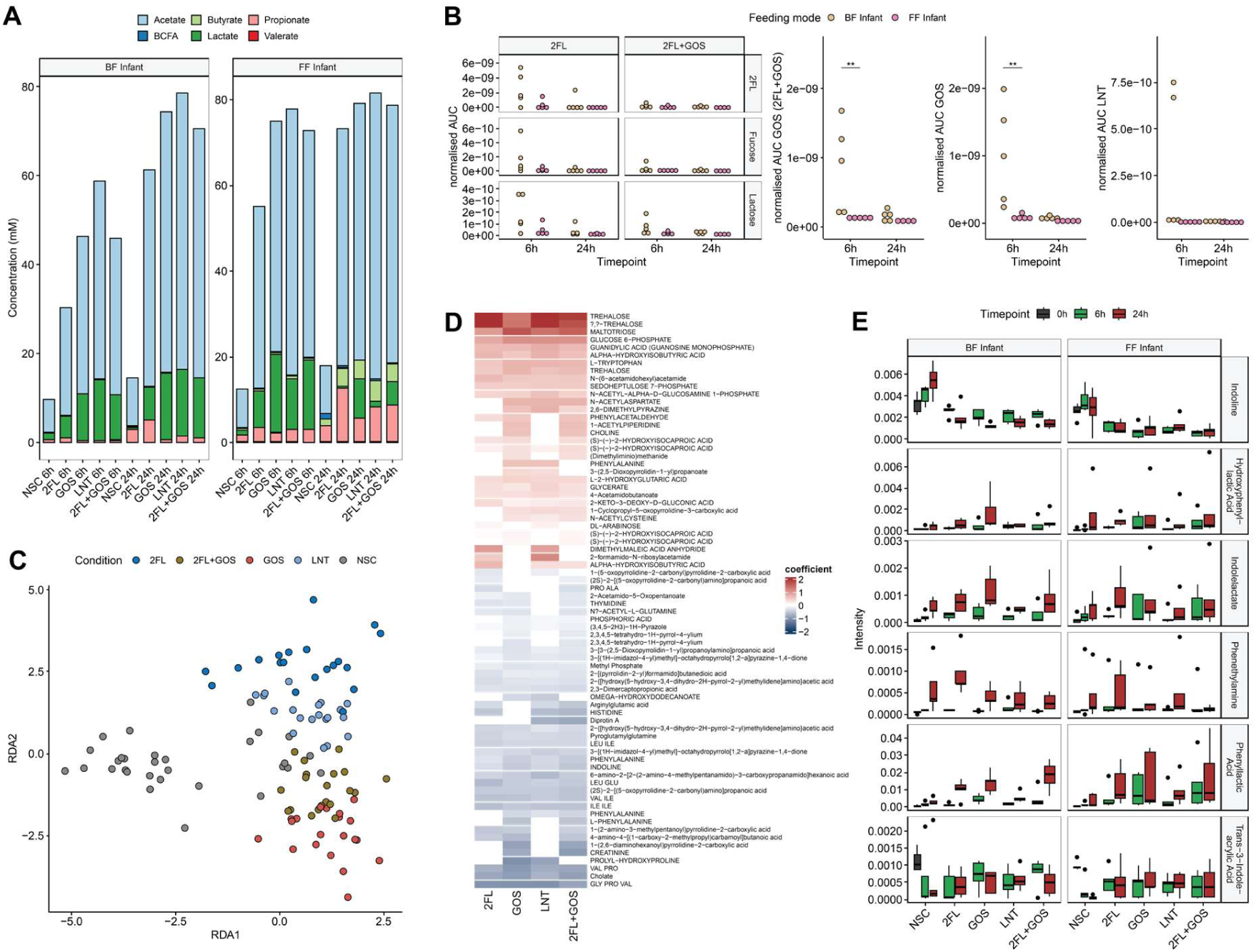
Prebiotic supplementation reorganizes carbon flux and amino acid metabolism in infant gut communities. (**A**) Short-chain fatty acid (SCFA) profiles across treatments reveal feeding-mode–dependent differences. FF communities show rapid lactate conversion and higher propionate and butyrate levels, whereas BF communities accumulate acetate and lactate. Prebiotics increase total SCFA production and reduce branched-chain fatty acids relative to non-substrate controls (NSC). (**B**) Prebiotic consumption over time, measured by HPAEC-PAD, demonstrates complete utilization by 24 h. FF communities metabolize GOS and LNT faster than BF communities. 2’FL supplementation transiently releases free lactose and, in some donors, fucose at 6 h, both depleted by 24 h, indicating substrate-specific generation of cross-feeding intermediates. (**C**) Partial treatment-constrained redundancy analysis (RDA) of untargeted metabolomics data shows distinct clustering of supplemented versus non-substrate control samples, by time- and prebiotic condition, reflecting substrate-driven rewiring of community metabolism. The 2′FL+GOS condition appears intermediate between the individual substrates. (**D**) Heatmap of metabolites significantly altered oner time (6 h vs 24 h) by prebiotic supplementation and culture time (log-fold change relative to NSC), showing increased carbohydrate-derived intermediates and reduced amino acid–related metabolites, consistent with a shift toward carbohydrate-driven energy metabolism. (**E**) Relative abundance of aromatic amino-acid derived metabolites, show substrate-, and feeding-mode–dependent profiles.

Substrate consumption analysis (HPEAC-PAD) showed complete utilization of prebiotics by 24h, with slower consumption in BF microbial community (**Figure 2B**). 2’FL supplementation transiently released free lactose and, in some donors, free fucose at 6h, and both were depleted by 24h. These intermediates likely serve as cross-feeding substrates for non-HMO-degrading taxa, supporting substrate-dependent metabolic interactions.

Untargeted polar metabolomics revealed distinct clustering by timepoint and supplementation condition (both p<0.001, PERMANOVA**; Figure 2C, Supplementary Figure S2B**). Differential analysis normalized to NSC showed consistent depletion of amino acid–derived metabolites under supplementation (**Figure 2D**, **Supplementary Figure S2C**). Hydroxy carboxylic acids, including hydroxy-isocaproic acid, were enriched under supplementation with minimal feeding-mode dependence (**Supplementary Figure S2D)**. Phenyllactic acid (PLA), derived from tyrosine, increased at 24 h under all prebiotic conditions except LNT, independent of the feeding mode, while phenylethylamine (PEA) was higher in BF donors, and modestly increased by 2’FL in both feeding-mode groups (**Figure 2E**). Hydroxyphenyllactic acid (HPLA) showed similar trends as PLA, however was less pronounced. In contrast, tryptophan-derived metabolites such as indolelactic acid (ILA) showed minimal changes across substrates, while indoline levels decreased under supplementation (**Figure 2E**).

Metagenomic mapping linked PLA/ILA and HPLA, which is derived from phenylalanine via the same enzymatic reactions, production primarily to lactic acid bacteria and Bifidobacteria (**Supplementary Excel Sheet**). Together, these data indicate that prebiotic supplementation redirects metabolic activity from amino acid fermentation toward carbohydrate-driven fermentation, while 2′FL uniquely enhances propionate production across feeding-mode-defined ecological states.

### 2’FL rewires cross-feeding interactions and metabolic dependencies in a defined microbial community

To investigate the mechanisms underlying the prebiotic effects on the microbial community, we constructed a six-member simplified defined microbial community (referred to as SynCom; **see Methods**) comprising *Phocaeicola vulgatus* (sensitive to prebiotic-induced inhibition), *Bifidobacterium breve* and *B. infantis* (dominant early-life species), *E. coli*, *Lactobacillus gasseri* and *Ruminococcus gnavus* (prevalent in early-life gut microbiota). The microbial taxa were selected based on their presence in our infant faecal samples **(Figure 1)**, and support from metagenomics and computational modelling studies (Alessandri et al., 2022; Casaburi et al., 2021; Versluis et al., 2022).

Individual species metabolic profiling revealed that both Bifidobacteria and *R. gnavus* primarily produced acetate and formate, with minor succinate, while Bifidobacteria and *L. gasseri* predominately secreted lactate after 48h of culture (**Figure 3A**, **Supplementary Figure S3A**). *P. vulgatus* and *E. coli* generated succinate, with minor propionate production. Only Bifidobacteria consumed LNT, and GOS with *B. breve* targeting lower-DP GOS, and *B. infantis* using broader fractions (**Figure 3B**, **Supplementary Figure S3B**). 2’FL was metabolized slightly by *B. infantis* and *P. vulgatus*, while *R. gnavus* cleaved 2’FL extracellularly into lactose and fucose (**Figure 3B, Supplementary Figure S3C**), providing substrates for cross-feeding. Cross feeding assays confirmed that *E. coli* consumed formate (**Figure 3C**), while *E. coli*, Bifidobacteria, and to a lesser extent, *P. vulgatus* consumed lactose and fucose (**Figure 3B**); *R. gnavus* consumed only fucose.

**Figure 3.**
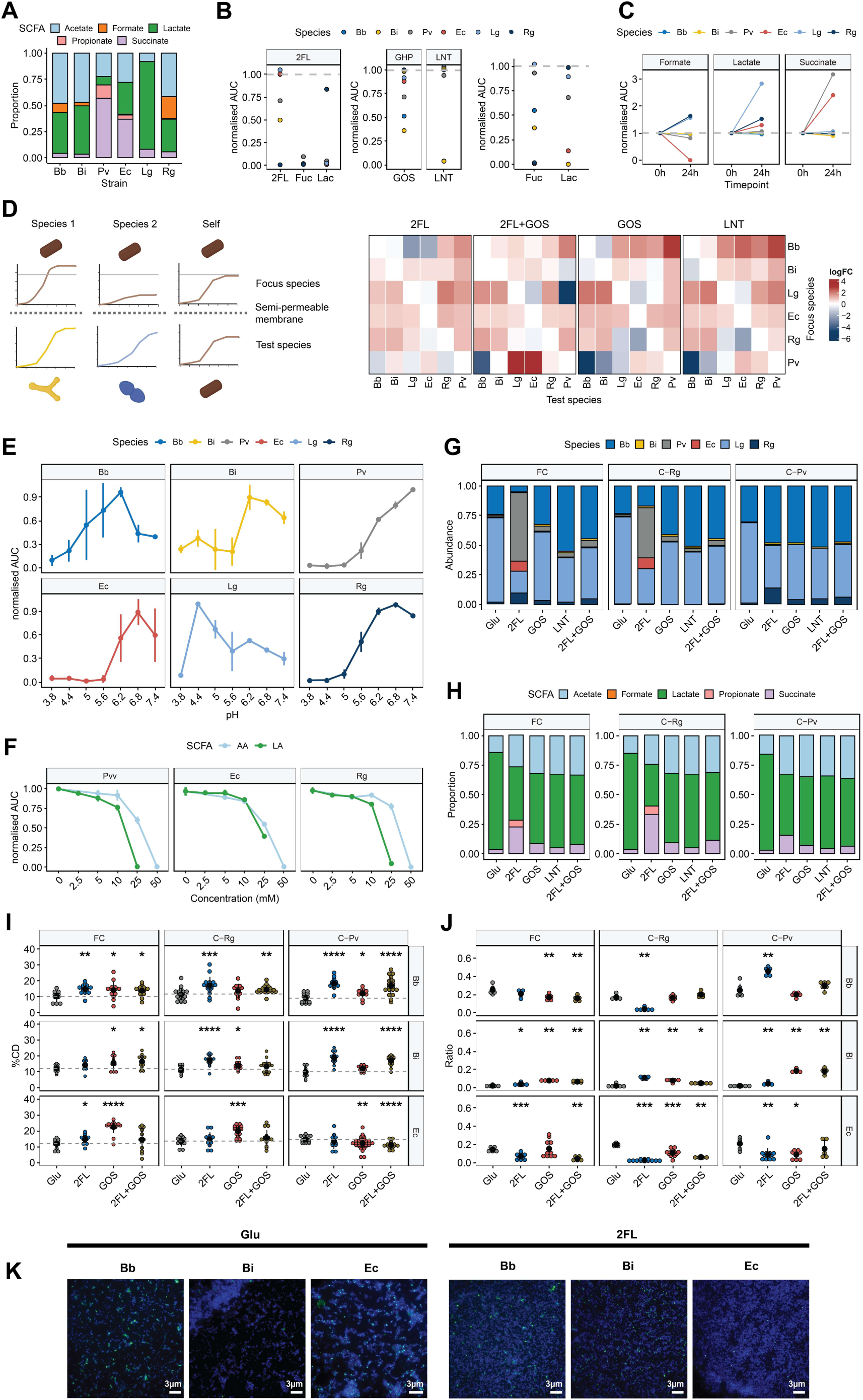
A synthetic infant gut community reveals extensive cross-feeding and species-specific interactions. (**A**) Average short-chain fatty acid (SCFA) production by each SynCom member highlights distinct metabolic profiles. (**B**) Prebiotic consumption normalised to initial substrate concentration. 2’FL is cleaved extracellularly, transiently releasing fucose and lactose. Right panel shows utilization of these monosaccharides by individual members, revealing substrate-specific preferences. (**C**) *E. coli* consumes formate, while any other major SCFAs are not metabolised by the remaining SynCom members, indicating selective cross-feeding pathways. (**D**) Binary co-culturing experiments reveal species interaction networks. Log-fold changes are calculated relative to each species’ monoculture (“self”) growth, highlighting inhibitory and facilitative effects among community members. (**E**) pH sensitivity of each SynCom member measured by its area under the curve of their respective growth curves, showing preference of Bb, Bi and Lg for lower pH (**F**) Area under the curve of growth curves when either acetic acid (AA) or lactic acid (LA) are added. This shows higher sensitivity of Pv and Rg towards lactate, compared to Ec (**G**) Relative community composition after 24h of culture across prebiotic supplementation. 2’FL supplementation selectively increases *P. vulgatus* and *E. coli* abundances. (**H**) SCFA composition of different community variants (full community (FC); communities lacking *R. gnavus* (C-Rg), or *P. vulgatus*, (C-Pv). 2’FL consistently increases succinate levels, while propionate and succinate production depends on the presence of *P. vulgatus* in FC and C-Rg communities. (**I**) Relative heavy water labelling efficiency as a proxy for metabolic activity. Prebiotic supplementation enhances metabolic activity across the community, with *E. coli* showing notably higher activity when *P. vulgatus* is present, illustrating interspecies metabolic dependencies. (**J**) Relative species abundances determined by FISH for the same culture as in **(I).** (**K**) Fluorescence in situ hybridisation (FISH) shows a string decrease in Bb and Ec abundance when 2’FL is supplemented and Rg is missing; Abbreviations: Bb – *B. breve*, Bi – *B. infantis*, Lg – *L. gasseri*, Ec – *E. coli*, Rg – *R. gnavus,* Pv – *P. vulgatus*.

Binary co-culture experiments with semipermeable membranes (Longhi et al., 2025) demonstrated that most species gained a growth advantage when paired, and that 2’FL uniquely altered interaction patterns compared to GOS or LNT, which displayed similar profiles (**Figure 3D**). Notably, inhibition of *P. vulgatus* by Bifidobacteria or *L. gasseri* was alleviated in the presence of 2’FL, illustrating a substrate-mediated mitigation of competitive suppression. As Bacteroides/*Phocaeicola* species are pH sensitive (Duncan et al., 2009; Keitel et al., 2024), we investigated *P. vulgatus* tolerance and found complete growth inhibition at pH 5, whereas Bifidobacteria and *L. gasseri* tolerated lower pH (**Figure 3E**). Surprisingly, *E. coli* and *R. gnavus*, both lactate producers, showed similar inhibition at these pHs, but were not inhibited in cross-feeding cultures with Bifidobacteria. This suggested that the type of secreted SCFA modulates inhibition. Indeed, the inhibition of *P. vulgatus* and *R. gnavus* was stronger by lactate than acetate at comparable concentrations, whereas *E. coli* was less affected (**Figure 3F**). Together these findings show an additive effect of pH and SCFA type on species interactions. The alleviation of *P. vulgatus* inhibition by 2’FL was pH independent, as binary cultures with GOS showed similar pH values (4.97 or 5.0 for *B. breve* and *B. infantis,* respectively), while 2’FL cultures showed slightly higher pH without affecting sensitivity (5.5 for *B. breve* or 4.97 for *B. infantis*) (**Supplementary Figure S3D**).

Community-level dynamics using a full community (FC) and two partial communities lacking either *P. vulgatus* (C^-Pv^) or *R. gnavus* (C^-Rg^) revealed mechanistic dependencies. 2’FL supplementation increased abundances of *P*. *vulgatus*, *R. gnavus*, and *E. coli* (**Figure 3G**), with *E. coli* showing strong dependence on *P. vulgatus* presence. Omission of *R. gnavus* boosted *B. breve* and *L. gasseri* abundance, while omission of *P. vulgatus* enhanced *R. gnavus*, indicating that nutrient competition and cross-feeding jointly dictate community composition. Lactate remained the dominant SCFA in all communities, while formate was depleted, likely through consumption by *E. coli* (**Figure 3H**). 2’FL-induced succinate and propionate production was primarily attributed to *P. vulgatus*, as shown by its absence in the C^-Pv^ community.

Single-cell activity levels determined by heavy water labelling, Raman microspectroscopy and fluorescens in situ hybridisation (FISH) (Berry et al., 2015), confirmed enhanced metabolic activity in Bifidobacteria, particularly in the presence of 2’FL (**Figure 3I**). Cross-feeding dependencies were also observed in *E. coli* in the presence of GOS, which displayed significantly higher activity when *P. vulgatus* was present. This dynamic effect appears to be growth-phase dependent or part of a larger cross-feeding network involving more species, as *E. coli* growth in *P. vulgatus* conditioned supernatants did not differ between GOS or glucose supplementation (**Supplementary Figure S3E**). FISH analysis further confirmed that *B. breve* and *E. coli* relied on *R. gnavus* in the 2’FL condition, likely through lactose and fucose cross-feeding, while *B. infantis* could invade the niche when *B. breve* was absent (**Figure 3J,K**). Collectively, the results mechanistically demonstrate that that prebiotic supplementation rewires species-specific cross-feeding networks, with *P. vulgatus* and *R. gnavus* acting as ecological hubs whose metabolism cascades through the community, which, in turn reshape downstream species interactions and metabolic outputs.

### Complex infant faecal communities recapitulate synthetic community interactions and metabolite patterns

Analysis of donor-derived faecal cultures revealed matches for SynCom members: *B. breve*, *B. infantis* (low prevalence), *E. coli,* and *R. gnavus*. *L. gasseri* was absent, but replaced by *Lactobacillus rhamnosus*, and *P. vulgatus* was not detected; instead, the closely related *P. dorei* was the predominant *Phocaeicola* species. Metagenomic bin analysis indicated that these strains were slightly phylogenetically distinct form the SynCom strains, but closely resembled them functionally with respect to their metabolic modules (**Supplementary Figure S4A-E**). This supports that mechanism elucidated in the SynCom are translatable to complex infant gut communities.

After 24h, feeding-mode-dependent differences in community composition emerged: *E. coli* was more abundant in BF-derived samples, and *P. dorei* dominated in FF-derived samples (**Figure 4A**). Pairwise Spearman correlations of species log-fold changes (relative to NSC) demonstrated strong negative correlations between *P. dorei* and both *Bifidobacterium* species in BF donors, consistent with the competitive or exclusionary interactions observed in the SynCom. These effects were attenuated in the presence of 2’FL and were not observed in FF communities (**Figure 4B**), highlighting the role of initial community structure in shaping prebiotic responses. Data from an independent infant cohort confirmed similar ecological signatures, with negative correlations between *Bacteroides/Phocaeicola* and *Bifidobacterium,* supporting the broader relevance of these interaction patterns (Casaburi et al., 2021)(**Supplementary Figure S4F**). Propionate levels correlated positively with *Bacteroides*/*Phocaeicola* abundance (**Figure 4C**), reinforcing their role as propionate producers in the SynCom. Together, these results demonstrate that key mechanistic features uncovered in the SynCom-Bifidobacterium–Bacteroides antagonism, feeding-mode-specific interaction patterns, and hub-linked metabolite production are recapitulated in complex infant faecal communities, underscoring the predictive value of simplified, hub-focused systems.

**Figure 4.**
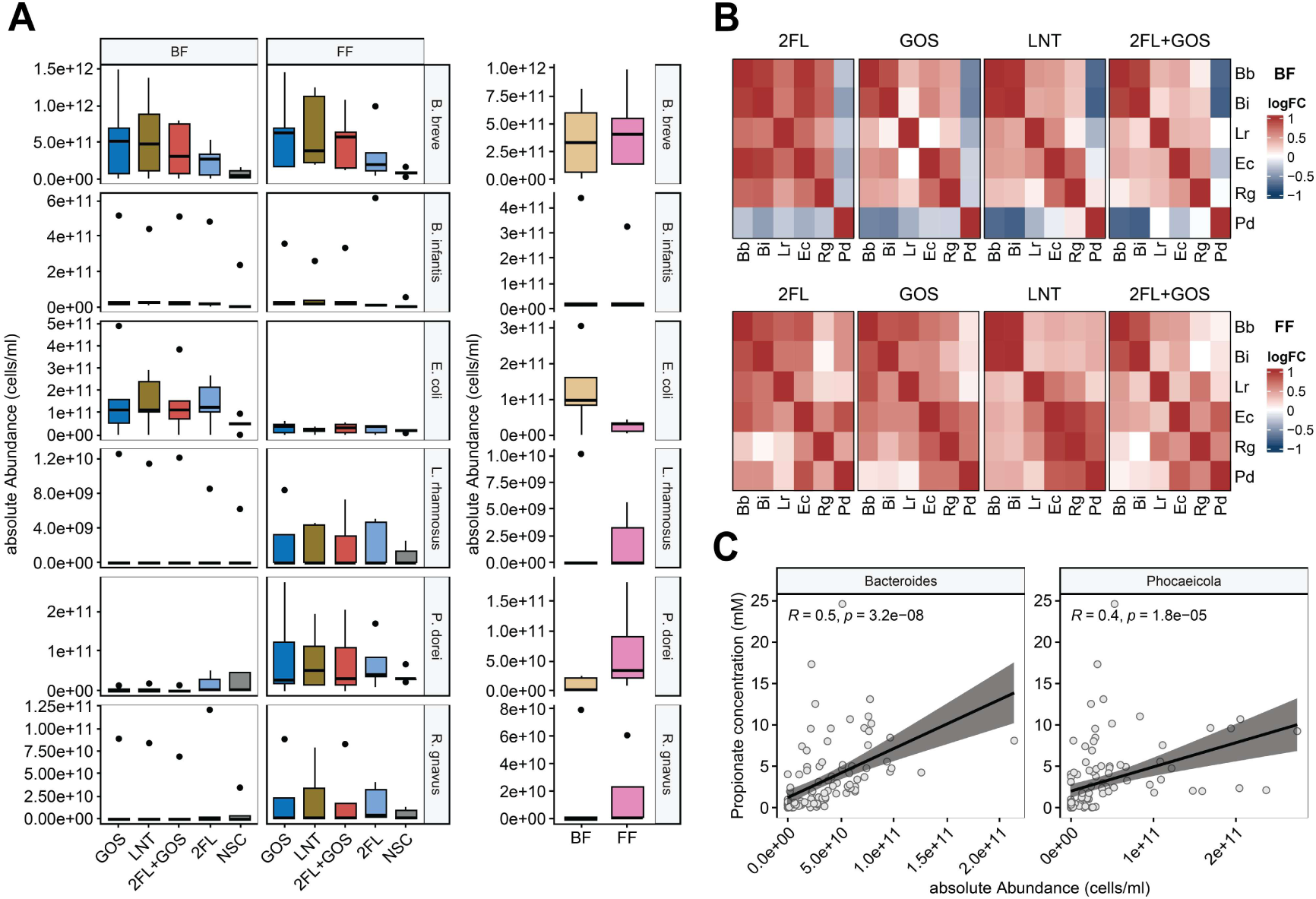
Key interactions inferred from the synthetic community are recapitulated in donor-derived faecal cultures. (**A**) Relative abundance of SynCom members or their closest relatives in donor faecal cultures after 24h. *P. vulgatus* was not detetced, but its close member *P. dorei* was present. Similarly, *L. gasseri* was absent, whereas *L. rhamnosus* was detected. Data are normalized to initial inoculum. (**B**) Heatmaps showing pairwise correlations of species abundances (log-fold change relative to non-substrate control) across donors, revealing patterns of competition and co-occurrence that reflect SynCom-derived interactions. (**C**) Propionate concentrations correlate significantly with the abundance of *Bacteroides* and *Phocaeicola* species, highlighting their role in community-level SCFA production. Abbreviations: Bb – *B. breve*, Bi – *B. infantis*, Ec – *E. coli*, Rg – *R. gnavus*, Lr – *L. rhamnosus*, Pd – *P. dorei*.

### Infant microbiome-associated microbial metabolites reversibly suppress colonic peristaltic activity ex vivo

Given that SCFAs detected in the SynCom **(Figure 2H)** are known modulators of intestinal motility, we hypothesised that SynCom-derived metabolites may alter colonic contractile activity. As *P. vulgatus* emerged as a hub species **(Figure 1E, 2G, J)**, we further investigated whether spent media from the full community and a *P. vulgatus*-deficient community exert differential effects using the ex vivo IntestiFlow motility platform (**Figure 5A, Supplementary text)**.

**Figure 5.**
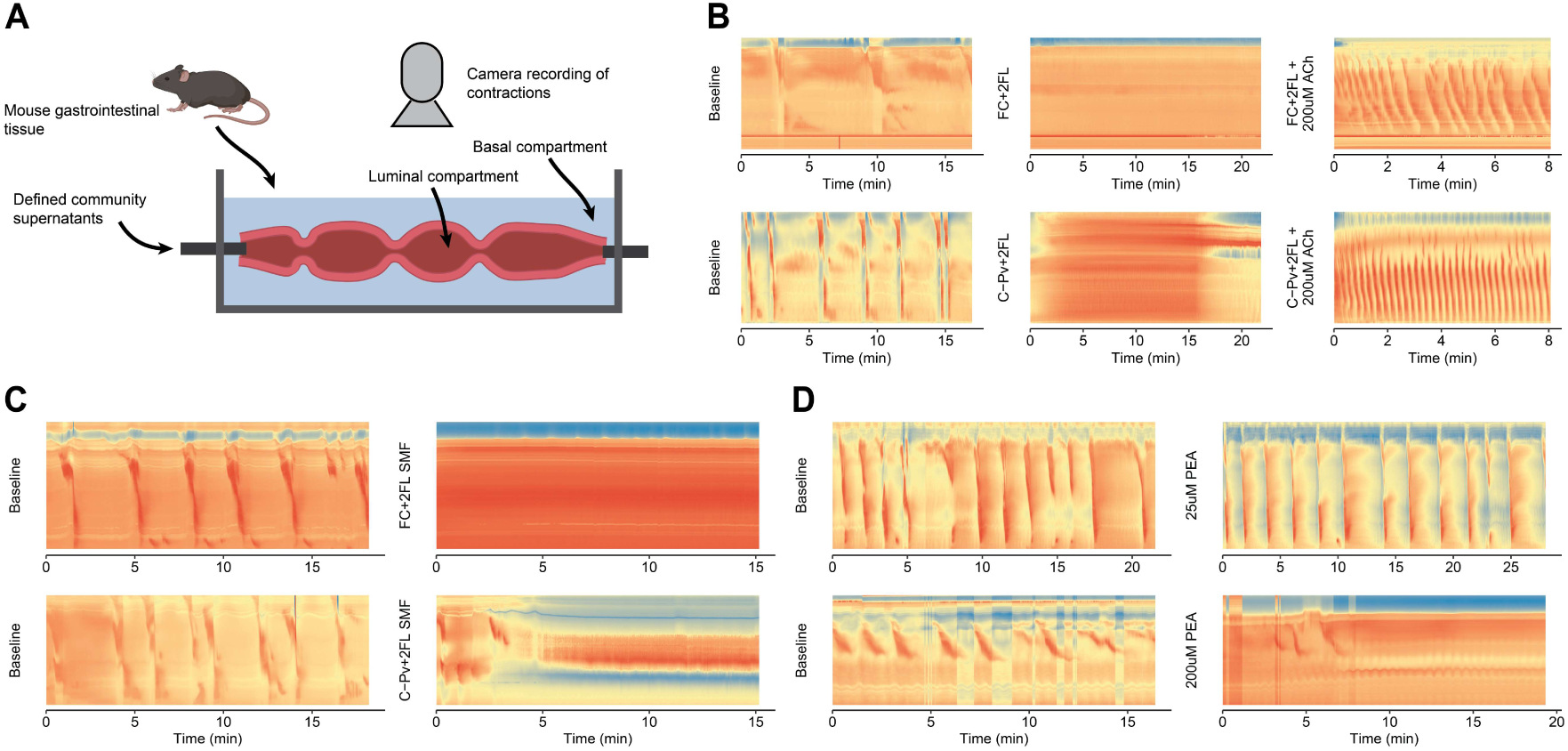
Infant gut–associated synthetic community metabolites reversibly suppress colonic contractility ex vivo. (**A**) Schematic of the IntestiFlow platform. Mouse colonic tissue is mounted in a perfusion chambre and connected to inlet and outlet tubing, allowing controlled administration of microbial supernatants and metabolites while contractile activity is recorded via time-laps imaging for spatiotemporal analysis. (**B**) Representative spatiotemporal diameter maps (**see Extended Methods**) illustrating colonic motility patterns. Supernatants from either the full community or the *P. vulgatus*-deficient community induce a marked inhibition in contractile activity, which is restored upon administration of 200 uM acetylcholine. Red signifies reduced luminal diameter (contraction), whereas blue/yellow indicates relaxation. (**C**) Methanolic extracts constituted of mostly small molecules (SMF) from the supernatants administered in (**A**) show similar inhibition of colonic contractility, as illustrated by representative spatiotemporal diameter maps. (**D**) Exposure to 200 µM, but not 25 µM, PEA induced tonic contraction, characterized by sustained elevation of basal tone and complete suppression of both peristaltic and segmental movements.

To test this, surplus mouse colonic tissue was perfused luminally under constant pressure with physiological buffer and contractile activity was quantified by video recording. Under baseline conditions, tissues exhibited an average of 0.3 full peristaltic contractions per minute over ∼20 minutes, after which perfusion was switched to treatment media **(Figure 5B, Supplementary Video S1)**. Administration of pH-adjusted spent media from either community resulted in a marked suppression of full peristaltic contractions. In the absence of peristaltic activity, tissues remained in a semi-contracted state. This inhibitory effect was reversible, as addition of 200 µM acetylcholine to the basal compartment restored robust peristaltic activity, increasing both contraction frequency (2.3 contractions per minute) and completeness relative to baseline.

To determine whether SCFAs were responsible for this phenotype, we applied synthetic SCFA mixtures matching the concentrations and composition detected in the SynCom **(Figure 2H)**. However, SCFA treatment alone did not reproduce the observed changes in motility **(Supplementary Figure S5A)**, suggesting that other microbially derived metabolites are responsible for the effect. To test this, we used methanolic extracts of spent community media, which capture a broad range of small molecules present in these supernatants. Again, we observed strong inhibition of colonic contractility, with no difference between extracts derived from either the full community or the *P. vulgatus*-deficient community **(Figure 5C)**, suggesting that small polar metabolites may underlie the observed effects.

One candidate molecule is phenethylamine, which has previously been reported to exert constrictive effects on intestinal smooth muscle (Batista-Lima et al., 2019), however was not tested on colonic tissue yet. We therefore investigated PEA as a candidate bioactive metabolite, as its production was found to be increased in our donor-derived faecal culture samples, especially when 2’FL was supplemented (**Figure 2E**). These observations were confirmed in single cultures of *R. gnavus,* one member of the SynCom, in a minimal culture medium supplemented with tyrosine and phenylalanine (**Supplementary Figure S5B**). Luminal administration of 200uM, but not 25uM PEA, indeed led to a persistent tonic contraction of the tissue, which was characterized by a persistent increase in basal colonic tone that fully suppressed both peristaltic and segmented movements, representing a strong inhibition of normal phasic motility (**Figure 5D**). The effect was dose-dependent, with an EC_50_ between 50-75uM, after correcting for interindividual variability in baseline contractility (**Supplementary Figure S5D**). Yet, these finding do not fully explain our observations as production of PEA in the SynCom was below the detection limit (data not shown), suggesting that other molecules are involved. Together, these results identify a previously unrecognised capacity of metabolites produced by an infant gut-associated synthetic community to robustly and reversibly suppress colonic motility. While neither SCFAs nor *P. vulgatus* were sufficient to explain this phenotype, PEA partially reproduced key aspects of the observed response, although only at concentrations exceeding those detected in the SynCom supernatant, suggesting that biogenic amines and additional community-derived metabolites may contribute to host neuromuscular modulation.

## Discussion

Prebiotic responses in the infant gut are shaped by interaction-dependent metabolic specialization rather than uniform bifidogenic expansion. Our findings suggest that dietary substrates reconfigure microbial community function through network-level rewiring of metabolic interactions, leading to microbial baseline-dependent community outputs.

Donor-derived fermentations recapitulated known compositional differences between BF and FF infants (**Figure 1B**) (Casaburi et al., 2021; Forbes et al., 2018; Ho et al., 2018; Ma et al., 2020), supporting the physiological relevance of our fermentation model. Prebiotics, including HMOs and GOS, consistently promoted *Bifidobacterium* growth across all donors and feeding modes, in line with previous reports (Bajic et al., 2020; Borewicz et al., 2019; Lindner et al., 2023). In contrast, *Bacteroides/Phocaeicola* species showed selective and donor-dependent responses, with growth inhibition primarily observed in BF-derived communities and alleviated by 2′FL supplementation. These findings indicate that prebiotic responsiveness is shaped by baseline community structure and manifests rapidly, within 6 h of culture. Importantly, these early effects occurred independently of detectable pH shifts or SCFA accumulation, suggesting that substrate-specific metabolism and interspecies interaction dynamics, including cross-feeding on yet unidentified metabolites, contribute to the observed responses beyond bulk acidification. Nevertheless, suppression of *Bacteroides* species is not consistently observed *in vivo* for infants receiving GOS containing formula (Barnett et al., 2023), highlighting that additional host- and diet-associated factors likely modulate these microbial interactions in natural settings.

Mechanistic dissection using a defined microbial community revealed extensive cross-feeding and substrate-dependent metabolic interdependencies **(Figure 3)**. Central to this network was the extracellular cleavage of 2’FL by *R. gnavus*, which expanded substrate availability, and supported downstream cross-feeding interactions, including promotion of *B. breve* growth via shared carbohydrate utilization pathways (Lou et al., 2023)(Seki et al., 2026). Consistent with previous reports (Boronat & Aguilar, 1981; Sawers, 2025), *E. coli* metabolised multiple fermentation intermediates, including formate, fucose, and lactose, highlighting its role as a flexible metabolic responder within the community.

A key emergent feature of the system was the identification of *P. vulgatus* as a functional ecological hub. Stable isotope probing and Raman-FISH analysis revealed that *E. coli* increased metabolic activity in the presence of *P. vulgatus* and GOS without corresponding changes in cell abundance, indicating that functional activity can be decoupled from compositional shifts. Such dissociation between metabolic activity and abundance has been described in other ecosystems, including marine microbial communities (Campbell et al., 2011) and suggests that *P. vulgatus* modulates the metabolic environment rather than simply promoting growth of specific taxa. Consistent with this role, *P. vulgatus* contributed substantially to succinate and propionate production, thereby shaping downstream metabolite availability within the ecosystem (**Figure 3A,H**) (Fernández-Veledo & Vendrell, 2019). Notably, similar hub-like behaviour, *P. vulgatus* has been shown in adult synthetic communities, where *P. vulgatus* can reprogram nutrient-driven interactions and promote metabolic adaptation in *Clostridioides difficile*, suggesting that this functional role may represent a conserved ecological feature rather than an infant-specific property (Sulaiman et al., 2025). Importantly, these interaction patterns were not recapitulated in monoculture, underscoring that metabolic behaviour emerges from community-level organisation rather than intrinsic properties of individual strains.

Our data further indicate that competitive dynamics within these communities are not governed solely by environmental acidification, but are strongly influenced by the chemical identity of fermentation products. Although several species exhibited comparable pH sensitivities, lactate exerted stronger inhibitory effects than acetate at similar concentrations, particularly on *P. vulgatus* and *R. gnavus* **(Figure 3E, F)**, indicating that metabolite-specific toxicity rather than bulk acidification shapes competitive outcomes. In contrast, lactate-producing species such as *E*.

*coli* displayed greater tolerance, suggesting that metabolite phenotype contributes to differential sensitivity to fermentation end products (Fang et al., 2022). Analogously, alleviation of *P. vulgatus* inhibition by 2’FL occurred independent of bulk pH differences, further supporting the notion that substrate-driven interaction rewiring, rather than acidification alone, determines community structure. While SCFA-mediated regulation of microbial growth has been extensively described in the context of pathogen control (Chang et al., 2024; Zhan et al., 2022), our findings extend this principle to commensal and hub taxa, highlighting that specific fermentation products act as selective ecological filters within the community.

We further found that metabolites produced by members of the SynCom induced a strong but reversible suppression of colonic contractility *ex vivo*. This suppression of peristaltic activity was accompanied by a sustained semi-contracted state, suggesting that microbial metabolites do not inhibit tissue activity, but alter the coordination between tonic and phasic motor programs. Such effects may arise through modulation of enteric neuromuscular signalling pathways controlling smooth muscle excitability and coordinated propagating contractions. Notably, SCFA mixtures matching SynCom metabolite profiles did not reproduce this phenotype, whereas methanolic extracts containing a broader spectrum of small nicrobial metabolites induced similar suppression of contractility. These findings suggest that other polar small microbial metabolites contribute to the observed effects. One candidate is PEA, which partially reproduced key aspects of the response, although inly at concentrations exceeding those detected in the SynCom supernatant, suggesting that PEA alone is unlikely to fully account for the observed phenotype. Instead, the effects may arise from the combined activity of multiple community-derived metabolites within the small polar fraction, potentially acting synergistically on enteric neuromuscular pathways. Together, these findings support the idea that the observed effects arise from a broader metabolite milieu rather than a single dominant compound. Mechanistically, microbial metabolites may interfere with cholinergic signalling pathways, including muscarinic receptor activity or downstream ion channel function, as previously suggested for other microbial products (Waclawiková et al., 2022). Functionally, altered peristaltic activity could influence intestinal transit and mixing dynamics, thereby affecting nutrient processing and luminal exposure times during early life in a context-dependent manner. Interestingly, suppression of peristaltic activity was accompanied by a sustained semi-contracted state, suggesting that microbial metabolites may not simply inhibit tissue activity, but rather alter the coordination between tonic and phasic motor programs. Such changes could reflect modulation of enteric neuromuscular signalling pathways controlling smooth muscle excitability and coordinated propagating contractions.

Collectively, our findings support a model in which prebiotic responses in early-life microbial communities are shaped by interaction-dependent metabolic organisation, where specific taxa act as ecological hubs that disproportionately influence metabolite production and community function. Rather than exerting uniform effects across taxa, dietary substrates rewire microbial networks by altering cross-feeding relationships, competitive sensitivities, and metabolite-mediated interactions. This framework integrates species-specific metabolism and community-level dynamics with host intestinal neuromuscular function, providing a unified view of how early-life dietary inputs shape microbiome function. Future longitudinal *in vivo* studies will be required to determine how these network-level processes influence microbiome maturation and host health trajectories.

## Material and Methods

### Fermentation experiments

Faecal samples from five exclusively breast-fed (BF) and five exclusively formula-fed (FF) infants were collected parental informed consent under approval of the Ethics Committee of the University Hospital Ghent (ref. BC-09977). Exclusion criteria included antibiotic use within 30 days and prior NEC or gut surgery. The mean donor age was 4.9 months, with both sexes represented.

Individual bioreactors (Cryptobiotix, Ghent, Belgium) containing 5 mL of nutritional medium (M0017, Cryptobiotix, Ghent, Belgium) were inoculated with faecal samples and supplemented with 5g/L 2’-fucosyllactose, 5g/L galactooligosaccharides, 5g/L lacto-N-tetraose (all FrieslandCampina), 5g/L of a 1:4 mixture of 2FL and GOS, or plain medium as non- substrate control (NSC). Concentrations have been chosen based on previous studies (Bajic et al., 2023). Cultures were incubated at 37 °C for 24 h under anaerobic conditions, with sampling at0 hours (NSC only), 6h and 24h. At each timepoint, pH, gas production, and cell counts (flow cytometry) were measured. SCFAs (acetate, propionate, butyrate, valerate) and branched chain fatty acids (bCFAs: sum of isobutyrate, isovalerate and isocaproate) were quantified via gas-chromatography with flame ionisation detection and lactate via a high-throughput spectrophotometric assay.

### Untargeted metabolomics

For untargeted metabolomics (see **Extended Methods** for additional information), 50 ul of sample was mixed with 250 ul ice-cold methanol, sonicated in a chilled bath, followed by addition of 500 ul ice-cold chloroform, mixed for 2 min and incubated on ice for 10 min, and 200 ul cold water. After incubation on ice and centrifugation (1000 × g, 5 min, 4 °C), 400 ul of the upper (polar) and lower (apolar) phases were recovered into clean tubes. The polar phase was diluted with 600 ul water and lyophilized overnight; the apolar phase was dried under nitrogen downflow for 30 min. Samples were stored at -20°C until analysis. Reconstitution was in 200 ul acetonitrile/water (1:1 v/v) for polar metabolites and or 100 ul isopropanol/acetonitrile/water (4:3:1, v/v/v) for apolar metabolites. SPLASH II Lipidomix internal standard was added to apolar samples.

For polar metabolites 1-μl of sample was injected for non-targeted metabolomics analysis by an Ultimate 3000 UHPLC system (Thermo Scientific, Dreieich Germany) and separated on a BEH-Amide column (100 mm × 2.1 mm,1.7 μm particle size, Waters, Massachusetts, USA) using a A: 0.1% formic acid in water, B: 0.1% formic acid in acetonitrile gradient. From apolar metabolites 10 μl of sample was injected and separated using a CSH- C18 column (100 mm × 2.1 mm,1.7 μm particle size, Waters, Massachusetts, USA) with a A: 0.1% formic acid and 10 mM ammonium formate in 60% acetonitrile 40% water, B: 0.1% formic acid and 10 mM ammonium formate in 90% 2-propanol and 10% acetonitrile gradient. Commonly between the two fractions, eluting analytes were electrosprayed in positive and negative mode (two separate injections) and analyzed by a quadrupole time of flight mass spectrometer (tims TOF Pro, Bruker, Bremen Germany).

The resulting data were analyzed using Metaboscape 5.0 (Bruker Daltonics, Bremen, Germany) to perform data deconvolution, peak-picking, and alignment of m/z features. Features were annotated with various sources of fragmentation spectra, as well as with SIRIUS (v6.0.7). For further downstream analysis the statistical programming language R (v4.3.3) was used. For both metabolite fractions separately, blank contaminants were removed, when a metabolite was present in the blank samples at 3 times the intensity compared to the analyte samples. Similarly features with high variability in the QC samples were removed if their coefficient of variance was one standard deviation above their overall mean. We further applied a prevalence filter, such that a feature had to be present in at least 25% of the samples. PCA and RDA analysis were performed with packages factoMineR (v2.11) and vegan (v2.6).

### Metagenomic sequencing

DNA was extracted using a QIAamp PowerFecal Pro DNA kit (Qiagen, Hilden, Germany) following manufacturer’s instructions, with the exception of using a mini bead beater for the lysis step. Library preparation was carried out using the Novogene NGS DNA Library Prep Set (Cat No.PT004) with index codes assigned to each sample. Briefly, genomic DNA was randomly fragmented to size of 350bp. The DNA fragments were end-polished, A-tailed, ligated with adapters. Size selection was performed, followed by Rolling Circle Amplification. The resulting PCR products were purified using the AMPure XP system and analysed for size distribution using an Agilent 2100 Bioanalyzer (Agilent Technologies, CA, USA). Quantification was performed using real-time PCR. The library was then sequenced using the Novaseq X plus flow cell with PE150 strategy. All sequencing data is available under the following project accession PRJNA1463617.

### Bioinformatic analysis

Briefly, raw reads were trimmed with trimGalore, and host DNA was removed with Hocort21 in biobloom mode, yielding “clean reads”. The clean reads were subsequently assembled using both SPAdes22 in meta mode or MEGAHIT 23 in parallel. Assemblies were subjected to binning using BASALT.24 The resulting bins were dereplicated using dRep25, yielding a non-redundant bin set. Taxonomic annotation of bins was performed using GTDB-Tk.26 In parallel, gene prediction was carried out on all (redundant) bins using Prokka27 and the predicted proteins were aggregated into a non-redundant protein catalogue using MMSeqs2.28 Bin abundances were determined by mapping clean reads against the non-redundant bin set using Bowtie 229 and mapped reads per contig were obtained by SAMtools idxstat.30 These steps were implemented in a Nextflow pipeline31, with environment files showing respective tool versions available online (https://github.com/m4rku5-5/metagenomicsPipeline.git). Read counts were converted to pseudo TPM values using the formula: pTPM = mapped reads / (total bin length / 1000) / (Total reads in sample / 1000000).

To generate absolute abundances for each bin, pTPM values were multiplied by cell counts (determined by flow cytometry, see above) and these values were used for downstream analysis. Compositional analysis was performed using the statistical programming language R (v4.3.3), with the following packages: phyloseq (v1.46.0), microViz (v0.12.1), vegan (v2.6), ggpubr (v0.6.0) and tidyverse (v2.0.0). Differential abundance of features was assessed using Maaslin232 (v1.16.0), including the Donor individual as random effect.

### HPAEC-PAD

For determining carbohydrate degradation profiles, bacterial cultures were centrifuged at 15000g for 10min at 4°C and supernatant was diluted 1:100 into fresh MilliQ water. Samples were injected into a Dionex ICS6000 system (ThermoFisher) equipped with a Dionex CarboPac PA1 analytical column (4x250mm) with respective pre-column. Depending on the compounds analysed two gradients were used. For GOS: 0min 0% eluent B, 20min 25%, 23min 100%; for all other analytes: 10min 0% eluent B, 28min 20%, 31min 100%; eluent A: 0.1M sodium hydroxide, eluent B: 0.1M sodium hydroxide + 1M sodium acetate. Detection was performed with pulsed amperometric detection using a gold electrode and carbohydrate quad waveform. Data was analysed using Chromeleon 7.2.9

### Bacterial cultures and construction of SynCom

All bacteria were grown under anaerobic conditions (5% CO2, 5% H2, balanced with N2) using an anaerobic vinyl chamber (Coy Laboratory Products) at 37°C. As medium either eGAM (enhanced GAM; for all culturing if not indicated differently) or adjusted M9 minimal medium (for metabolic labelling and secondary metabolite production) was used (**Supplementary Excel Sheet**). Strain designations of all species and appropriate mixing ratios, based on baseline growth rates, for the synthetic community can be found in the **Supplementary Excel Sheet.** For setup of the SynCom we first determined baseline growth rates in eGAM medium containing only glucose, by diluting species 1:100 from a two day culture. In order to avoid overgrowth of fast growing species and to yield an abundance equalised community we then determined the N0 of each strain and calculated a N0 which would be needed to reach OD 0.2 at 5h of growth. The ratio between original and new N0 was then used as new dilution ratio. Before pooling the SynCom each individual 48h strain culture was centrifuged at 3000g for 10min at RT and resuspended in fresh medium.

### Determination of binary bacterial interactions

For determining bacterial interactions between all combinations of synthetic community members bacteria were grown for 48h in standard eGAM. We used the duet co-culture system (Cerillo, Charlottesville, VA, USA) to contain each species in a separate well, but allow for metabolites to pass freely between the two chambers. As one plate only holds 18 combinations, experiments had to be split across multiple runs. One “focus species” was inoculated into one side of the well and test species were placed in the other side. All bacteria were tested in a 1:50 dilution, with respective prebiotics added (4g/L final concentration). Plates were sealed with Breath-Easy plate seals (Sigma Aldrich) and growth was monitored at 620nm with a MultiSkan FC plate reader (ThermoFisher) for 48h at 37°C and medium shaking every 20min. To determine growth impacts, growth curves were fit using the growthcurver R package (v0.3.1) and carrying capacity k was used as output. We then normalised the growth of each focus species when cultured together with another species to its self-growth (same species both sides).

### Determination of bacterial metabolic activity

To measure metabolic activity the community was grown accordingly for 24h with respective prebiotic additions (4g/L) in eGAM. After, cultures were centrifuged for 10min at 3000g and RT until resuspension in adjusted M9 minimal medium containing the respective prebiotic and 50% heavy water. Cultures were diluted 1:7 and incubated for 24h, until 200ul were fixed with 1ml of 4% PFA for 2h. Samples were washed 2 times with PBS and then stored in the freezer until further use. To determine labelling and identity of cells fluorescence in situ hybridisation (FISH) and Raman spectroscopy was carried out. FISH was performed using a standard liquid FISH protocol as described previously (Ge et al., 2022) with either BlongInf, PBR2 or Gam42 probes (**Supplementary Excel Sheet**). For single-cell activity measurements, samples (2uL) were spotted on aluminum-coated slides (Al136; EMF Corporation) and washed by dipping into ice-cold Milli-Q (MQ) water to remove traces of buffer components. Raman spectroscopy was carried out using a Renishaw inVia confocal microscope with an 100x air objective, at 532nm, 1800 grating and 12.5% laser power. Only cells displaying fluorophore signal from FISH were measured. Bleaching of the fluorophore signal was enabled for 20s, before 25s of acquisition. Raw spectra were manually curated for outliers and baseline corrected. To express percentage of labelling (%CD) the area under the curve of each CD (2040-2300nm) and CH (2800-3100nm) peak was used.

To determine abundances of each targeted group, cells were observed under an Olympus IX85, using DAPI as a counter stain. We performed image segmentation on at least 3 images using ilastik (v1.4.0.post1) and Fiji (v1.54g), which were then expressed as FISH labelled cells per total cells in field of view.

### SCFA determination

Supernatant of bacterial cultures was collected by centrifugation at 20000g for 10min and 4°C. One volume of 1M sulfuric acid was added and particulates were removed by filtration. Samples were the injected into a Shimadzu HPLC system equipped with a Nexera X2 SPD-M30A detector and the RezexTM ROA-Organic Acid H+ (8%) 300 x 7.8 mm LC column. Samples were run with 5nM sulfuric acid at 0.5 ml/min in isocratic conditions.

### Community sequencing

DNA was extracted from 200ul of sample using repeated bead beating as described previously (Yu & Morrison, 2004). The full length 16S region was then amplified using 35F with overhang (3’-AGAGTTTGATCCTGGCTCAGGATGAACG) and generic 1492R primers using NEB LongAmp (New England Biolabs, Ipswich, USA) with 100ng of template DNA. Resulting amplicons were purified using AMPure XP beads (Beckmann Coulter, Brea, USA) and library preparation was carried out using a 16S Barcoding Kit (Oxford Nanopore, Oxford, United Kingdom). Libraries were sequenced on a Flongle flow cell (Oxford Nanopore, Oxford, United Kingdom) and resulting reads were analysed using the EPI2ME software in *minimap* mode with a custom database consisting of only reference 16S sequences from respective community members.

### Measurement of gut motility

For determination of motility we used an in-house organ bath system, IntestiFlow, which is described in more detail in **Extended Methods**. Briefly, both male and female surplus mice (umbrella project number AVD11100-2021-15219) are terminated by cervical dislocation, colonic tissue is excised and stored on ice-cold oxygenated Tyrode’s solution (pH 7.4) until further usage. Tissue is slowly warmed up, hyperfused to remove any remaining pellets and then transferred to the organ bath system. Culture supernatants were prepared by centrifugation and subsequent filtration through a 0.2 um filter and stored at -20 °C until further use. Methanolic extracts were generate by addition of 40ml of ice cold methanol to 10ml of pH adjusted culture supernatant, followed by incubation for 2h on ice. Samples were centrifuged at 10000g for 10min at 4 °C, supernatant was collected, evaporated under nitrogen and resuspended in Tyrode’s solution. Community supernatants, SCFA mixes or methanolic extracts were added luminally under constant pressure control after equilibration of tissues for several minutes and baseline contractility recording for about 20min. All conditions were tested on individual tissue sections to minimize carryover of effects which could arise from sequential testing. Contractions were observed by means of camera recordings and then transferred to spatiotemporal maps using custom Phython code. Contraction strength is expressed by dividing mean width of the tissue in relaxation state by mean value of contraction width.

## Acknowledgment

This project was a part of the Fascinating programme and has been made possible in part by a contribution from the National Program Groningen and Province of Groningen, as well as FrieslandCampina.

## References

Alessandri, G., Fontana, F., Mancabelli, L., Lugli, G. A., Tarracchini, C., Argentini, C., Longhi, G., Viappiani, A., Milani, C., Turroni, F., Van Sinderen, D., & Ventura, M. (2022). Exploring species-level infant gut bacterial biodiversity by meta-analysis and formulation of an optimized cultivation medium. Npj Biofilms and Microbiomes, 8(1), 88. 10.1038/s41522-022-00349-1

Azad, M. B., Bridgman, S. L., Becker, A. B., & Kozyrskyj, A. L. (2014). Infant antibiotic exposure and the development of childhood overweight and central adiposity. International Journal of Obesity, 38(10), 1290–1298. 10.1038/ijo.2014.119

Bajic, D., Niemann, A., Hillmer, A.-K., Mejias-Luque, R., Bluemel, S., Docampo, M., Funk, M. C., Tonin, E., Boutros, M., Schnabl, B., Busch, D. H., Miki, T., Schmid, R. M., van den Brink, M. R. M., Gerhard, M., & Stein-Thoeringer, C. K. (2020). Gut Microbiota-Derived Propionate Regulates the Expression of Reg3 Mucosal Lectins and Ameliorates Experimental Colitis in Mice. Journal of Crohn’s and Colitis, 14(10), 1462–1472. 10.1093/ecco-jcc/jjaa065

Bajic, D., Wiens, F., Wintergerst, E., Deyaert, S., Baudot, A., & Van Den Abbeele, P. (2023). HMOs Exert Marked Bifidogenic Effects on Children’s Gut Microbiota Ex Vivo, Due to Age-Related Bifidobacterium Species Composition. Nutrients, 15(7), 1701. 10.3390/nu15071701

Barnett, D. J. M., Endika, M. F., Klostermann, C. E., Gu, F., Thijs, C., Nauta, A., Schols, H. A., Smidt, H., Arts, I. C. W., & Penders, J. (2023). Human milk oligosaccharides, antimicrobial drugs, and the gut microbiota of term neonates: Observations from the KOALA birth cohort study. Gut Microbes, 15(1), 2164152. 10.1080/19490976.2022.2164152

Batista-Lima, F. J., Rodrigues, F. M. D. S., Gadelha, K. K. L., Oliveira, D. M. N. D., Carvalho, E. F., Oliveira, T. L., Nóbrega, F. C., Brito, T. S., & Magalhães, P. J. C. (2019). Dual excitatory and smooth muscle-relaxant effect of β-phenylethylamine on gastric fundus strips in rats. Clinical and Experimental Pharmacology and Physiology, *46*(1), 40–47. 10.1111/1440-1681.13033

Berry, D., Mader, E., Lee, T. K., Woebken, D., Wang, Y., Zhu, D., Palatinszky, M., Schintlmeister, A., Schmid, M. C., Hanson, B. T., Shterzer, N., Mizrahi, I., Rauch, I., Decker, T., Bocklitz, T., Popp, J., Gibson, C. M., Fowler, P. W., Huang, W. E., & Wagner, M. (2015). Tracking heavy water (D_2_ O) incorporation for identifying and sorting active microbial cells. Proceedings of the National Academy of Sciences, 112(2). 10.1073/pnas.1420406112

Borewicz, K., Suarez-Diez, M., Hechler, C., Beijers, R., de Weerth, C., Arts, I., Penders, J., Thijs, C., Nauta, A., Lindner, C., Van Leusen, E., Vaughan, E. E., & Smidt, H. (2019). The effect of prebiotic fortified infant formulas on microbiota composition and dynamics in early life. *Scientific Reports*, 9(1), 2434. 10.1038/s41598-018-38268-x

Boronat, A., & Aguilar, J. (1981). Metabolism of L-fucose and L-rhamnose in Escherichia coli: Differences in induction of propanediol oxidoreductase. Journal of Bacteriology, 147(1), 181–185. 10.1128/jb.147.1.181-185.1981

Campbell, B. J., Yu, L., Heidelberg, J. F., & Kirchman, D. L. (2011). Activity of abundant and rare bacteria in a coastal ocean. Proceedings of the National Academy of Sciences of the United States of America, 108(31), 12776–12781. 10.1073/pnas.1101405108

Casaburi, G., Duar, R. M., Brown, H., Mitchell, R. D., Kazi, S., Chew, S., Cagney, O., Flannery, R. L., Sylvester, K. G., Frese, S. A., Henrick, B. M., & Freeman, S. L. (2021). Metagenomic insights of the infant microbiome community structure and function across multiple sites in the United States. Scientific Reports, 11(1), 1472. 10.1038/s41598-020-80583-9

Chang, K. C., Nagarajan, N., & Gan, Y.-H. (2024). Short-chain fatty acids of various lengths differentially inhibit Klebsiella pneumoniae and Enterobacteriaceae species. *mSphere*, S(2), e00781-23. 10.1128/msphere.00781-23

Darabi, B., Rahmati, S., HafeziAhmadi, M. R., Badfar, G., & Azami, M. (2019). The association between caesarean section and childhood asthma: An updated systematic review and meta-analysis. *Allergy*, Asthma & Clinical Immunology, 15(1). 10.1186/s13223-019-0367-9

Duncan, S. H., Louis, P., Thomson, J. M., & Flint, H. J. (2009). The role of pH in determining the species composition of the human colonic microbiota. Environmental Microbiology, 11(8), 2112–2122. 10.1111/j.1462-2920.2009.01931.x

Fang, Y., Stanford, K., & Yang, X. (2022). Lactic Acid Resistance and Population Structure of Escherichia coli from Meat Processing Environment. Microbiology Spectrum, 10(5), e01352–22. 10.1128/spectrum.01352-22

Fernández-Veledo, S., & Vendrell, J. (2019). Gut microbiota-derived succinate: Friend or foe in human metabolic diseases? Reviews in Endocrine and Metabolic Disorders, 20(4), 439–447. 10.1007/s11154-019-09513-z

Forbes, J. D., Azad, M. B., Vehling, L., Tun, H. M., Konya, T. B., Guttman, D. S., Field, C. J., Lefebvre, D., Sears, M. R., Becker, A. B., Mandhane, P. J., Turvey, S. E., Moraes, T. J., Subbarao, P., Scott, J. A., Kozyrskyj, A. L., & for the Canadian Healthy Infant Longitudinal Development (CHILD) Study Investigators. (2018). Association of Exposure to Formula in the Hospital and Subsequent Infant Feeding Practices With Gut Microbiota and Risk of Overweight in the First Year of Life. JAMA Pediatrics, *172*(7), e181161. 10.1001/jamapediatrics.2018.1161

Ge, X., Pereira, F. C., Mitteregger, M., Berry, D., Zhang, M., Hausmann, B., Zhang, J., Schintlmeister, A., Wagner, M., & Cheng, J.-X. (2022). SRS-FISH: A high-throughput platform linking microbiome metabolism to identity at the single-cell level. Proceedings of the National Academy of Sciences, *119*(26), e2203519119. 10.1073/pnas.2203519119

Ho, N. T., Li, F., Lee-Sarwar, K. A., Tun, H. M., Brown, B. P., Pannaraj, P. S., Bender, J. M., Azad, M. B., Thompson, A. L., Weiss, S. T., Azcarate-Peril, M. A., Litonjua, A. A., Kozyrskyj, A. L., Jaspan, H. B., Aldrovandi, G. M., & Kuhn, L. (2018). Meta-analysis of effects of exclusive breastfeeding on infant gut microbiota across populations. *Nature Communications*, S(1), 4169. 10.1038/s41467-018-06473-x

Hoskinson, C., Dai, D. L. Y., Del Bel, K. L., Becker, A. B., Moraes, T. J., Mandhane, P. J., Finlay, B. B., Simons, E., Kozyrskyj, A. L., Azad, M. B., Subbarao, P., Petersen, C., & Turvey, S. E. (2023). Delayed gut microbiota maturation in the first year of life is a hallmark of pediatric allergic disease. Nature Communications, 14(1). 10.1038/s41467-023-40336-4

Keitel, L., Miebach, K., Rummel, L., Yordanov, S., & Büchs, J. (2024). Process analysis of the anaerobe Phocaeicola vulgatus in a shake flasks and fermenter reveals pH and product inhibition. Annals of Microbiology, 74(1), 7. 10.1186/s13213-023-01745-4

Lindner, C., Looijesteijn, E., Dijck, H. V., Bovee-Oudenhoven, I., Heerikhuisen, M., Broek, T. J. V. D., Marzorati, M., Triantis, V., & Nauta, A. (2023). Infant Fecal Fermentations with Galacto-Oligosaccharides and 2′-Fucosyllactose Show Differential Bifidobacterium longum Stimulation at Subspecies Level. Children, 10(3), 430. 10.3390/children10030430

Longhi, G., Petraro, S., Milani, C., Tarracchini, C., Argentini, C., Vergna, L. M., Lugli, G. A., Mancabelli, L., Lee, C., Turroni, F., van Sinderen, D., & Ventura, M. (2025). Genetic Characterisation of the upp Gene in Bifidobacterium bifidum PRL2010. Microbial Biotechnology, 18(7), e70189. 10.1111/1751-7915.70189

Lou, Y. C., Rubin, B. E., Schoelmerich, M. C., DiMarco, K. S., Borges, A. L., Rovinsky, R., Song, L., Doudna, J. A., & Banfield, J. F. (2023). Infant microbiome cultivation and metagenomic analysis reveal Bifidobacterium 2’-fucosyllactose utilization can be facilitated by coexisting species. Nature Communications, 14, 7417. 10.1038/s41467-023-43279-y

Ma, J., Li, Z., Zhang, W., Zhang, C., Zhang, Y., Mei, H., Zhuo, N., Wang, H., Wang, L., & Wu, D. (2020). Comparison of gut microbiota in exclusively breast-fed and formula-fed babies: A study of 91 term infants. Scientific Reports, 10(1), 15792. 10.1038/s41598-020-72635-x

Milani, C., Duranti, S., Bottacini, F., Casey, E., Turroni, F., Mahony, J., Belzer, C., Delgado Palacio, S., Arboleya Montes, S., Mancabelli, L., Lugli, G. A., Rodriguez, J. M., Bode, L., de Vos, W., Gueimonde, M., Margolles, A., van Sinderen, D., & Ventura, M. (2017). The First Microbial Colonizers of the Human Gut: Composition, Activities, and Health Implications of the Infant Gut Microbiota. Microbiology and Molecular Biology Reviews, 81(4). 10.1128/mmbr.00036-17

Milani, C., Duranti, S., Bottacini, F., Casey, E., Turroni, F., Mahony, J., Belzer, C., Delgado Palacio, S., Arboleya Montes, S., Mancabelli, L., Lugli, G. A., Rodriguez, J. M., Bode, L., De Vos, W., Gueimonde, M., Margolles, A., Van Sinderen, D., & Ventura, M. (2017). The First Microbial Colonizers of the Human Gut: Composition, Activities, and Health Implications of the Infant Gut Microbiota. Microbiology and Molecular Biology Reviews, 81(4). 10.1128/mmbr.00036-17

Needham, B. D., Funabashi, M., Adame, M. D., Wang, Z., Boktor, J. C., Haney, J., Wu, W.-L., Rabut, C., Ladinsky, M. S., Hwang, S.-J., Guo, Y., Zhu, Q., Griffiths, J. A., Knight, R., Bjorkman, P. J., Shapiro, M. G., Geschwind, D. H., Holschneider, D. P., Fischbach, M. A., & Mazmanian, S. K. (2022). A gut-derived metabolite alters brain activity and anxiety behaviour in mice. Nature, 602(7898), 647–653. 10.1038/s41586-022-04396-8

Sawers, R. G. (2025). How FocA facilitates fermentation and respiration of formate by Escherichia coli. Journal of Bacteriology, 207(2), e00502–24. 10.1128/jb.00502-24

Seki, D., Pollak, S., Kujawska, M., Kiu, R., Acuna-Gonzalez, A., Crouch, L. I., Bakshani, C. R., Chivers, P. T., Mommers, M., van Best, N., Penders, J., & Hall, L. J. (2026). Human milk oligosaccharide mediates mutualism between Escherichia coli and Bifidobacterium bifidum. Nature Communications, 17(1), 3489. 10.1038/s41467-026-71764-7

Shao, Y., Forster, S. C., Tsaliki, E., Vervier, K., Strang, A., Simpson, N., Kumar, N., Stares, M. D., Rodger, A., Brocklehurst, P., Field, N., & Lawley, T. D. (2019). Stunted microbiota and opportunistic pathogen colonization in caesarean-section birth. Nature, 574(7776), 117–121. 10.1038/s41586-019-1560-1

Soyyılmaz, B., Mikš, M. H., Röhrig, C. H., Matwiejuk, M., Meszaros-Matwiejuk, A., & Vigsnæs, L. K. (2021). The Mean of Milk: A Review of Human Milk Oligosaccharide Concentrations throughout Lactation. Nutrients, 13(8), 2737. 10.3390/nu13082737

Sulaiman, J. E., Thompson, J., Cheung, P. L. K., Qian, Y., Mill, J., James, I., Im, H., Vivas, E. I., Simcox, J., & Venturelli, O. S. (2025). Phocaeicola vulgatus shapes the long-term growth dynamics and evolutionary adaptations of Clostridioides difficile. Cell Host & Microbe, 33(1), 42–58.e10. 10.1016/j.chom.2024.12.001

Tsukuda, N., Yahagi, K., Hara, T., Watanabe, Y., Matsumoto, H., Mori, H., Higashi, K., Tsuji, H., Matsumoto, S., Kurokawa, K., & Matsuki, T. (2021). Key bacterial taxa and metabolic pathways affecting gut short-chain fatty acid profiles in early life. The ISME Journal, 15(9), 2574–2590. 10.1038/s41396-021-00937-7

Van Den Abbeele, P., Deyaert, S., Thabuis, C., Perreau, C., Bajic, D., Wintergerst, E., Joossens, M., Firrman, J., Walsh, D., & Baudot, A. (2023). Bridging preclinical and clinical gut microbiota research using the ex vivo SIFR® technology. Frontiers in Microbiology, 14, 1131662. 10.3389/fmicb.2023.1131662

Versluis, D. M., Schoemaker, R., Looijesteijn, E., Muysken, D., Jeurink, P. V., Paques, M., Geurts, J. M. W., & Merks, R. M. H. (2022). A Multiscale Spatiotemporal Model Including a Switch from Aerobic to Anaerobic Metabolism Reproduces Succession in the Early Infant Gut Microbiota. mSystems, 7(5), e00446–22. 10.1128/msystems.00446-22

Waclawiková, B., Codutti, A., Alim, K., & El Aidy, S. (2022). Gut microbiota-motility interregulation: Insights from *in vivo, ex vivo* and *in silico* studies. Gut Microbes, 14(1), 1997296. 10.1080/19490976.2021.1997296

Yu, Z., & Morrison, M. (2004). Improved extraction of PCR-quality community DNA from digesta and fecal samples. BioTechniques, *36*(5), 808–812. 10.2144/04365ST04

Zhan, Z., Tang, H., Zhang, Y., Huang, X., & Xu, M. (2022). Potential of gut-derived short-chain fatty acids to control enteric pathogens. Frontiers in Microbiology, 13. 10.3389/fmicb.2022.976406

