## Supplementary Text for "Feeding-mode-defined microbial communities modulate prebiotic responses and alters colonic motility in early life"

### Extended methods

### Description of the ex vivo IntestiFlow Platform

*System Setup*

The IntestiFlow platform is a low-cost, pressure-controlled, perfused ex vivo system for continuous measurement of intestinal motility, designed as an alternative to classical ring-based organ bath systems (**Figure 0A-B**). The core of the system is a 3D-printed PLA culture reservoir, which can accommodate up to four tissue segments and is positioned on a custom heating pad to maintain physiological temperatures. PTFE tubing is used as standard tubing throughout the system and serves as the inlet/outlet ports for perfusion and attachment points for securing the tissue lumen.

To achieve precise and stable control of luminal pressure, we employed pressure-driven flow instead of syringe pumps. Piezoelectric micropumps (Bartels Mikrolectronik, Germany) pressurize the headspace of sealed falcon tubes containing the perfusate, resulting in controlled displacement of liquid into the tissue lumen.. Intraluminal pressure is monitored continuously using in-line pressure sensors (Honeywell, USA), and maintain ed at a constant maintained at a constant setpoint via a Python-based PID controller running on a BananaPi Zero. Maintaining stable luminal pressure is critical, as pressure strongly influences intestinal motility patterns (Schreiber, 2014). Video recordings are acquired using a Full HD camera with custom LED illumination.

*Image analysis*

Image analysis is carried out using custom Python code. Raw video frames undergo the following steps (**Figure 0C, Supplementary Video S1**): 1- Selection of the region corresponding to the culture reservoir; 2- Application of SLIC super-pixel segmentation (*skimage*) using three segments (empirically determined to best separate lumen and tissue); 3- Counting the number of pixels corresponding to the segment capturing the contracting tissue wall, column-wise along the gut axis; 4- Storage of the resulting 1D signal (time × gut length) as arrays for further processing.

Spatiotemporal motility maps are generated after temporal down sampling using *ggplot* in R.

*Tissue preparation and Measurement of motility*

Prior to experiments the system is assembled, and all tubing is purged. Oxygenated Tyrode's solution (pH 7.4) is used for both luminal and basal compartments. Mice are euthanized by cervical dislocation, and the whole colon is excised from the anus. Tissue is immediately transferred to cold, oxygenated Tyrode's solution; optimal motility is observed where transfer times are kept under 10-20 minutes.

Tissue is gradually warmed to 37°C over ~ 5 minutes and mounted in the system by securing the inlet with surgical thread, ensuring no air is introduced into the lumen. Lumen is gently perfused to remove fecal pellets. Afterwards, the reservoir is rinsed, and refilled with fresh buffer, and the tissue is fully connected to both in- and outlet ports. Video recording and PID-controlled perfusion are started once the absence of air bubbles is confirmed.

Motility recordings are consistently stable for 2-3 hours, although measurements are limited to ≤ 2 hours to prevent confounding by tissue fatigue. To validate the system, basolateral administration of acetylcholine induced strong phasic contraction, confirming functional neuromuscular responsiveness (**Figure 0D**). Tissue viability and structural integrity were further assessed histologically, demonstrating preservation of tissue architecture over the duration of motility experiments (**Figure 0E**).

*Resource availability*

All model files and code are available under https://github.com/m4rku5-5/IntestiFlow

**Untargeted metabolomics**

For untargeted metabolomics, 50 ul of sample was mixed with 250 ul ice-cold methanol, sonicated in a chilled bath, followed by addition of 500 ul ice-cold chloroform , mixed for 2 minutes and incubated on ice for 10 minutes, and 200 ul cold water. After incubation on ice and centrifugation (1000 × g, 5 min, 4 °C), 400 ul of the upper (polar) and lower (apolar) phases were recovered into clean tubes. The polar phase was diluted with 600 ul water and lyophilized overnight; the apolar phase was dried under nitrogen downflow for 30 min. Samples were stored at -20°C until analysis. Reconstitution was in 200 ul acetonitrile/water (1:1 v/v) for polar metabolites and or 100 ul isopropanol/acetonitrile/water (4:3:1, v/v/v) for apolar metabolites. SPLASH II Lipidomix internal standard was added to apolar samples.

For polar metabolites 1-µl of sample was injected for non-targeted metabolomics analysis by an Ultimate 3000 UHPLC system (Thermo Scientific, Dreieich Germany). Using a binary solvent system (A: 0.1% formic acid in water, B: 0.1% formic acid in acetonitrile) metabolites were separated on a BEH-Amide column (100 mm × 2.1 mm,1.7 µm particle size, Waters, Massachusetts, USA) with a VanGuard 2.1 mm × 5 mm guard column of the same material, by applying a linear gradient from 99% B to 40% B in 6 minutes and then to 4% B in 2 minutes at a flow rate of 0.4 ml/min, returning to initial conditions in 0.1 minute and equilibration for 2 minutes (10 minute total run time). From apolar metabolites 10 µl of sample was injected by an Ultimate 3000 UHPLC system (Thermo Scientific, Dreieich Germany). Using a binary solvent system (C: 0.1% formic acid and 10 mM ammonium formate in 60% acetonitrile 40% water, D: 0.1% formic acid and 10 mM ammonium formate in 90% 2-propanol and 10% acetonitrile) metabolites were separated on a CSH- C18 column (100 mm × 2.1 mm,1.7 µm particle size, Waters, Massachusetts, USA) with a VanGuard 2.1 mm × 5 mm guard column of the same material, by applying a linear gradient from 40% D to 43% D in 3 minutes then to 50% D in 0.1 minutes followed by a change to 54% D in 10 minutes then to 70% D in 0.1 minutes and to 99% D in 5.9 minutes holding at 99% D for 0.5 minutes before returning to initial conditions in 0.1 minutes and equilibration for 1 minute at a flow rate of 0.4 ml/min (20 minute total run time).

Commonly between the two fractions, eluting analytes were electrosprayed in positive and negative mode (two separate injections) by an Apollo II ion funnel ESI source (Bruker Daltonics Inc). Source settings were as follows: capillary voltage 4500 V or -3500 V; end plate offset 500 V; drying temperature 250 °C; desolvation gas (nitrogen) flow 8.0 l $min^{-1}$; nebulizer gas pressure 3.0 bar. Samples were analyzed by an quadrupole time of flight mass spectrometer (tims TOF Pro, Bruker, Bremen Germany) in times-off mode, auto MS/MS settings were: switching threshold 350 clear-to-send (cts); cycle time 0.5 s; active exclusion after 3 spectra; release after 0.2 min reconsidering precursors if ratio current/previous intensity > 2. Analytes selected for fragmentation were fragmented by collision with nitrogen gas at a collision energy of 20-30 eV. Precursors and fragments were analyzed by the time-of-flight analyzer using a range of 20-1300 m/z.

The resulting data were analyzed using Metaboscape 5.0 (Bruker Daltonics, Bremen, Germany) to perform data deconvolution, peak-picking, and alignment of m/z features using the T-ReX 3D peak extraction and alignment algorithm. All spectra were recalibrated on an internal lockmass segment (NaFormate clusters) and peaks were extracted with a minimal peak length of 8 spectra (7 for recursive extraction), and an intensity threshold of 1000 cts. The resulting feature tables contain deconvoluted features (adducts combined based on chromatographic correlation and known mass differences). Features were annotated, using SMARTFORMULA (narrow threshold, 1.5 ppm, mSigma:10; wide threshold 5.0 ppm, mSigma:30), to calculate a molecular formula. Spectral libraries including Bruker MetaboBASE 3.0, Bruker HDBM 2.0, MetaboBASE 2.0 in silico, MSDIAL LipidDBs, MoNA VF NPL QTOF, and GNPS, were used for feature annotation (narrow threshold, 1.5 ppm, mSigma 10, msms score 900, wide threshold 5.0 ppm, mSigma:30 msms score 800). Analyte lists of in house measured validated standards were also used to annotate features based on mass, retention time and fragmentation spectrum narrow threshold, 1.5 ppm, RT within 6 seconds, mSigma 10, msms score 900, wide threshold 5.0 ppm, RT within 12 seconds, mSigma:30 msms score 800). An annotated feature was considered of high confidence if more than two green boxes were present in the Annotation Quality column of the program and low confidence if less than two green boxes were present. We also identified metabolites by analysis with SIRIUS (v6.0.7), only retaining structure annotations with an exact confidence score greater than 0.5.

### Supplementary Figures

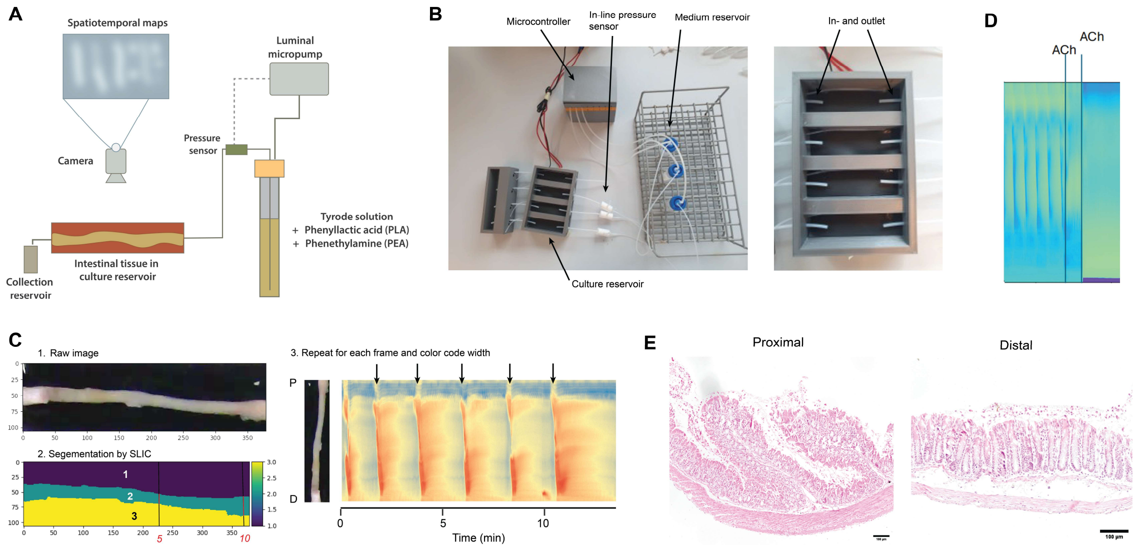


**Figure S0. Design and validation of the IntestiFlow platform for ex vivo gut motility measurements.** (**A**) Schematic overview of the IntestiFlow Platform, including micropumps, controller, pressure sensor, culture reservoir and camera. Micropumps pressurize the reservoir headspace, displacing liquid and driving luminal perfusion towards the mounted tissue. (**B**) Photograph of the assembled system (**C**) Schematic of spaciotemporal map (STMaps) generation. Raw video frames are segmented into three regions using the SLIC algorithm. For each pixel column, the number of pixels assigned to segment 2, corresponding to the radial width of the tissue, is counted. These width-values are color coded for each frame to generate an STMap, where the y-axis represents the proximal to distal axis of the tissue. Peristaltic contractions appear as transient reductions in width. (**D**) Basolateral acetylcholine administration induces strong tonic contractions, confirming neuromuscular responsiveness of the tissue. (**E**) Hematoxylin-Eosin-stained tissue sections from either proximal or distal colonic tissue show no apparent tissue damage after luminal perfusion for 2h.

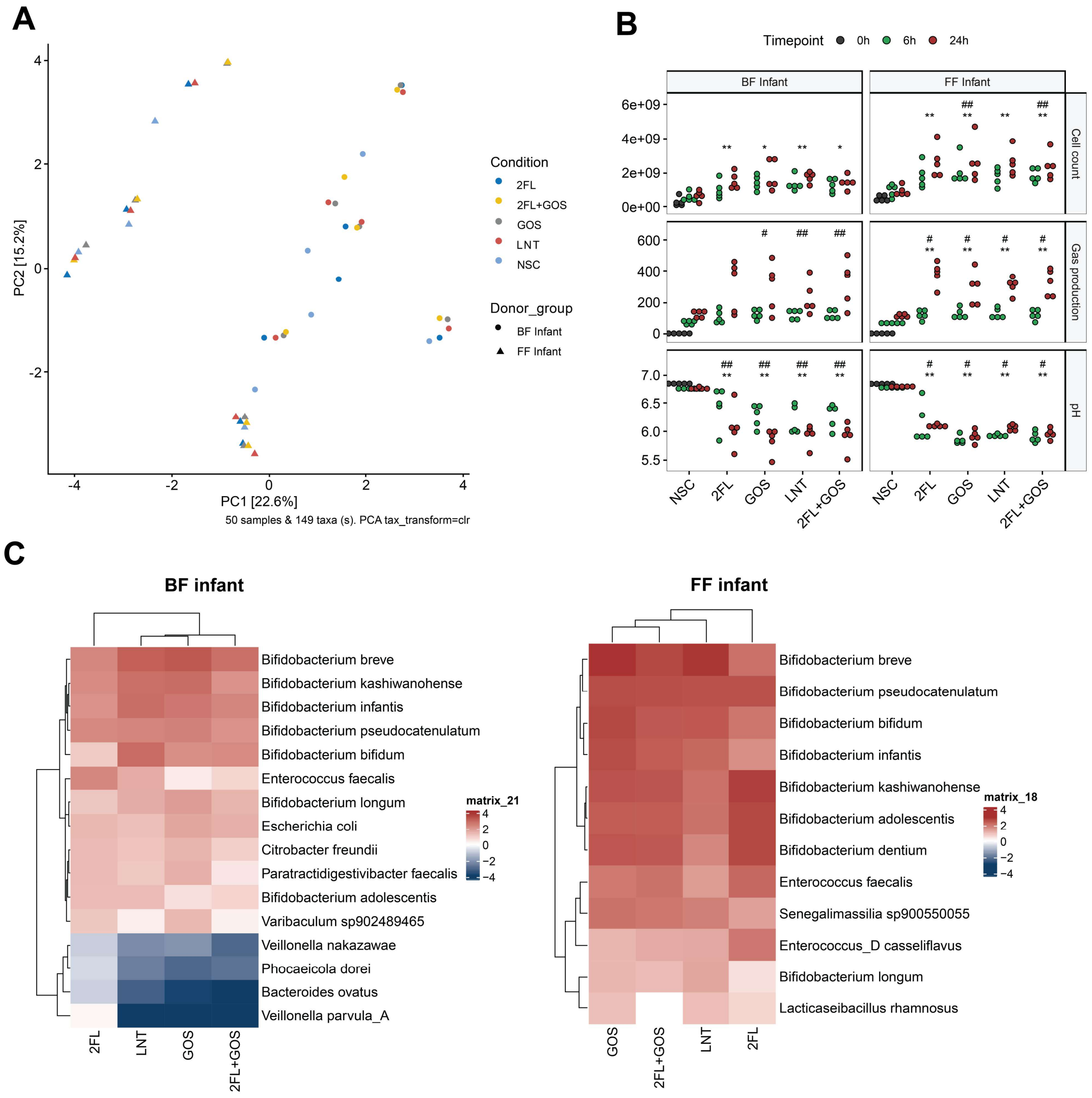


**Figure S1. Donor-specific effects on infant gut fermentation and species responses to prebiotics.** (**A**) Principal component analysis (PCA) showing strong clustering of samples by donor, indicating that donor identity overrides the effects of prebiotic supplementation. (**B**) Key fermentation parameters. Gas production did not differ between between feeding modes, but was increased by prebiotic supplementation, particularly at 24h. (**C**) Heatmap similar to **Figure1E**, stratified by feeding mode, showing inhibition of *Bacteroides/Phacaeicola* species occurs primarily in BF donor samples.

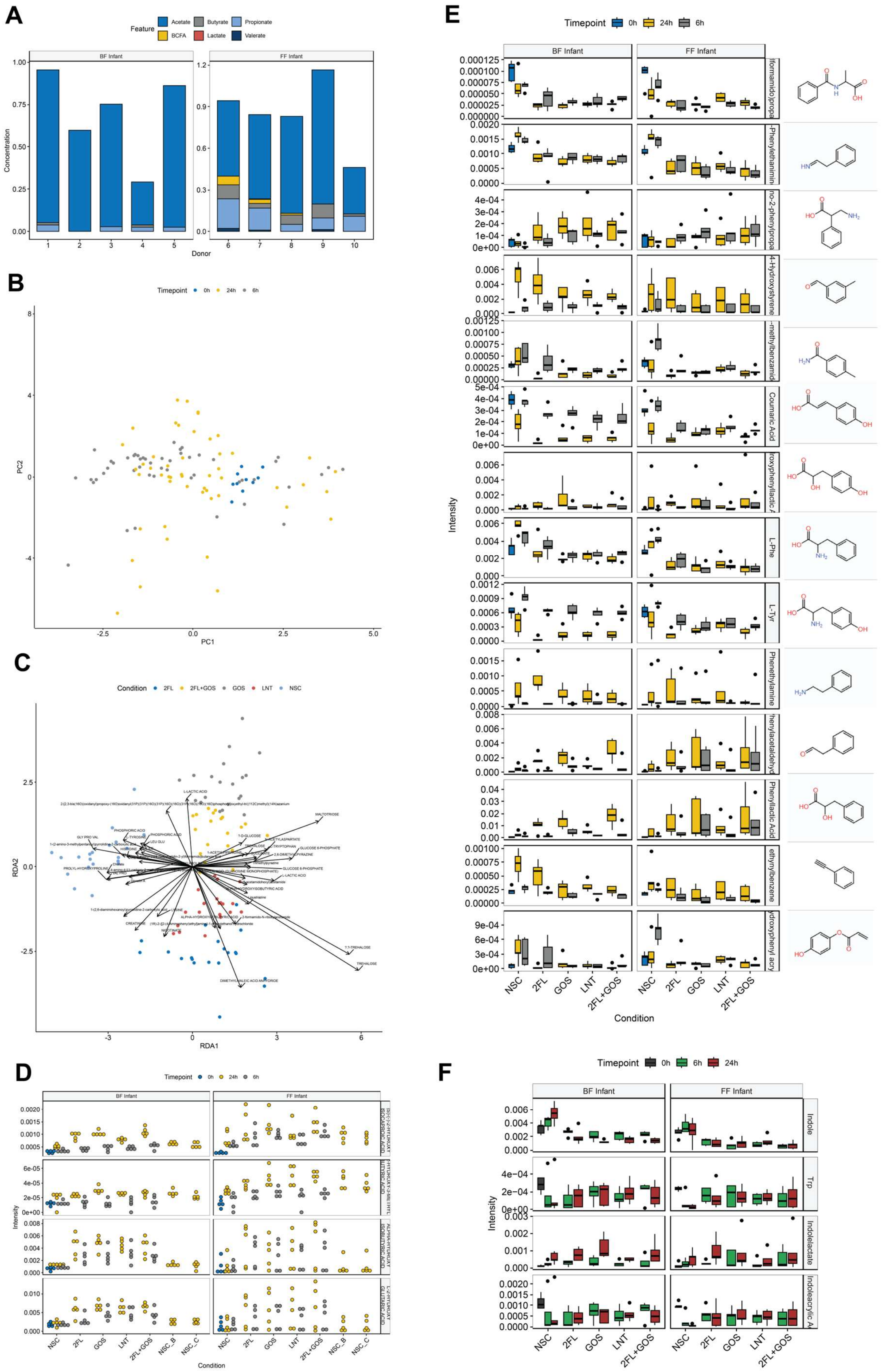

A
Feature
Acetate
Butyrate
Propionate
BCFA
Lactate
Valerate
BF Infant
FF Infant
Concentration
Donor
B
Timepoint
0h
24h
6h
PC1
PC2
C
Condition
2FL
2FL+GOS
GOS
LNT
NSC
RDA1
RDA2
L-LACTIC ACID
MALTOTRIOSE
TREHALOSE
DIMETHYLMALEIC ACID ANHYDRIDE
D
Timepoint
0h
24h
6h
BF Infant
FF Infant
Intensity
Condition
NSC
2FL
GOS
LNT
2FL+GOS
NSC_B
NSC_C
E
Timepoint
0h
24h
6h
BF Infant
FF Infant
Coumaric Acid
L-Phe
L-Tyr
Phenethylamine
Phenyllactic Acid
Intensity
Condition
F
Timepoint
0h
6h
24h
BF Infant
FF Infant
Indole
Trp
Indolelactate
Intensity

**Figure S2. Prebiotic supplementation alters metabolite profiles in infant gut fermentations.** (**A**) SCFA levels at 0h, recapitulating baseline differences between donors, including higher BCFA, propionate and butyrate abundance in FF donor samples. (**B**) PCA analysis of untargeted metabolomics data showing clustering primarily by timepoint. (**C**) Redundancy analysis (RDA) analysis, constrained by prebiotic treatment. Arrows represent loadings, with direction indicating association with specific clusters. Prebiotic supplementation reduced amino acid metabolism, as loadings for these compounds point toward non-substrate control (NSC) samples. (**D**) Abundance, represented and peak intensities, of selected hydroxy carboxylic acids that are differentially abundant across treatments. (**E**) Selected amino-acid derived compounds identified via structure matching of SIRIUS predicted structures.

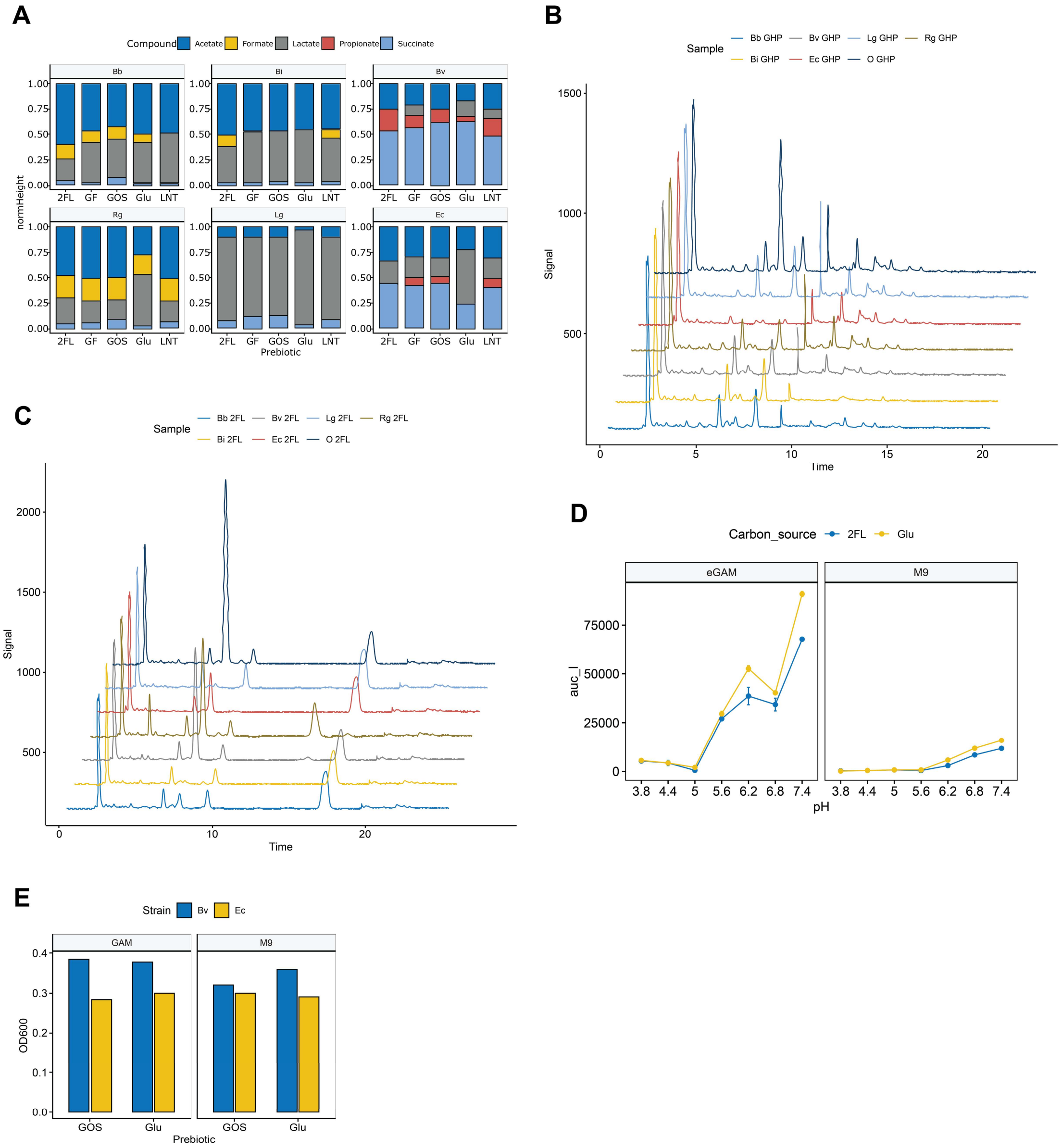


**Figure S3. Prebiotic utilization and short-chain fatty acid production by synthetic infant gut community members.** (**A**) SCFA profiles of each SynCom member under different prebiotic supplementation conditions. (**B**) HPAEC-PAD chromatograms showing differential GOS utilization patterns by *B. breve* and *B. infantis* (**C**) HPAEC-PAD chromatograms demonstrating release of free fucose and lactose when 2'FL is incubated with sterile *R. gnavus* culture supernatant, indicating extracellular fucosidase activity. (**D**) pH sensitivity of *P. vulgatus* in either eGAM or M9 medium, with either glucose or 2'FL as carbon source.

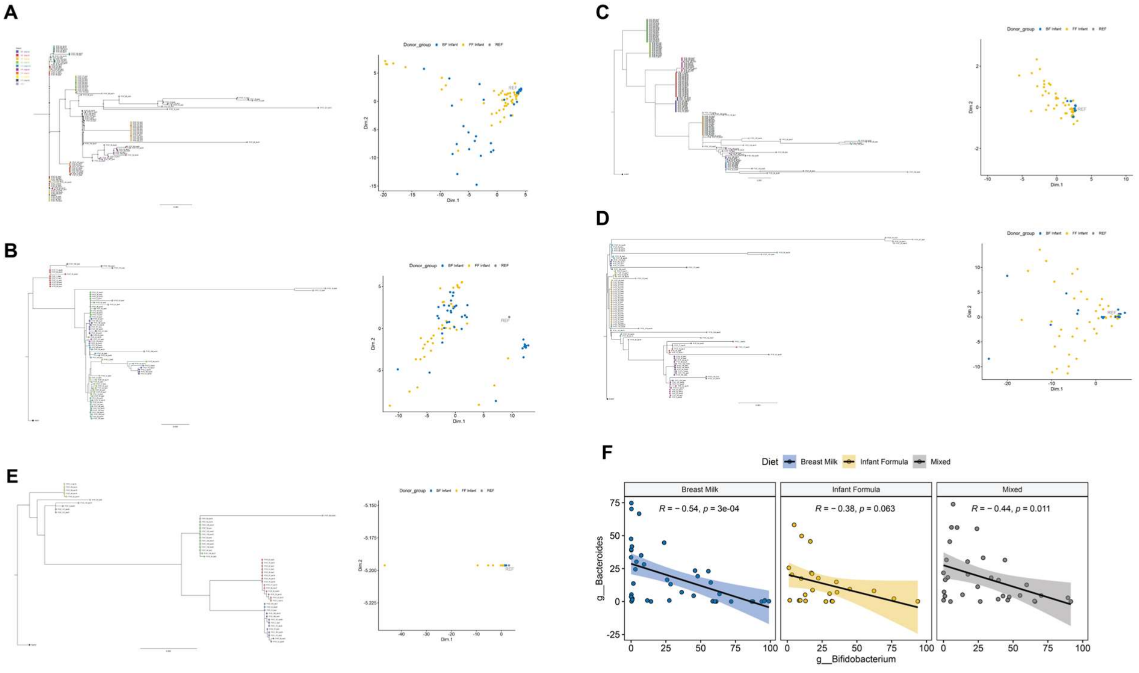


**Figure S4. Complex infant fecal samples recapitulate synthetic community interactions.** (**A**) Phylogenetic tree of all bins of *B. breve*, including the strain used in the SynCom (REF), as well as a PCA analysis of the metabolic modules. Similar analysis for *B. infantis* (**B**), *E. coli* (**C**), *P. vulgatus* (**D**), *R. gnavus* (**E**). (**F**) Correlation analysis between *Bifidobacterium* and *Bacteroides* abundances in (Casaburi et al., 2021).

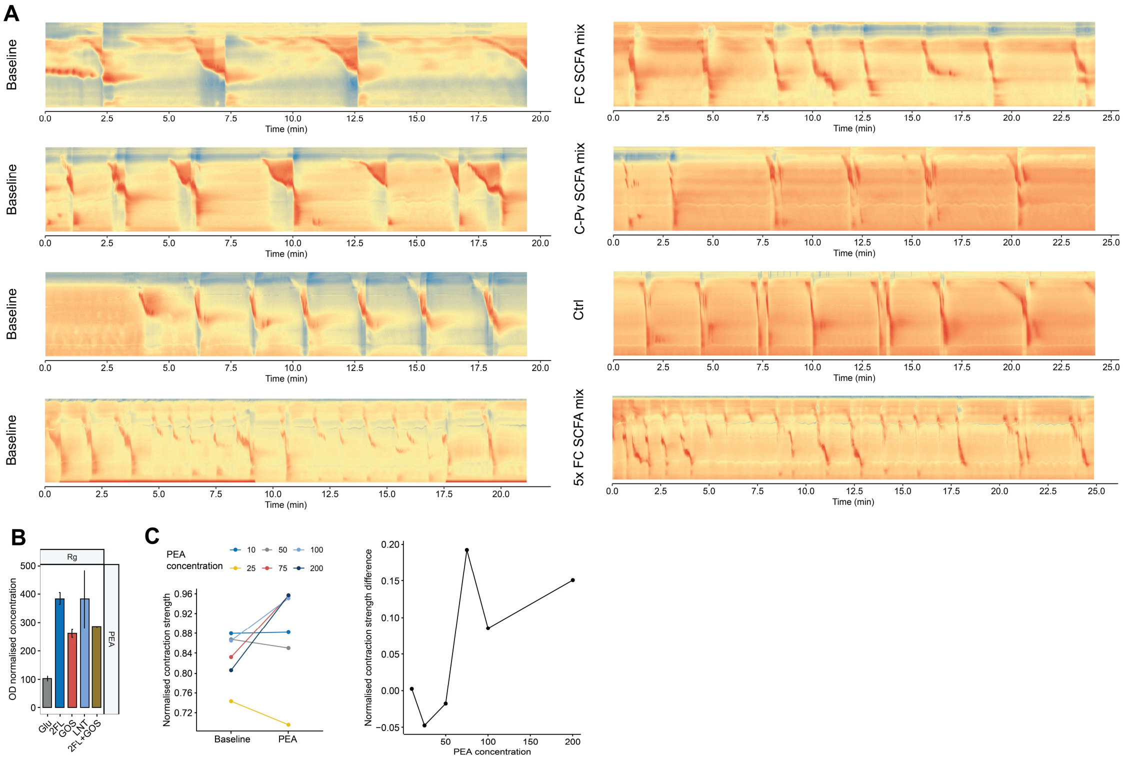


**Figure S5. Phenethylamine, but not short-chain fatty acids, modulates colonic contractility.** **(A)** Administration of a short-chain fatty acid (SCFA) mixture, formulated to match concentrations and composition observed in the synthetic community, did not induce substantial changes in colonic contraction patterns at either basal or fivefold elevated concentrations. The mixture was prepared in Tyrode's solution and adjusted to physiological pH using NaOH. To control for increased sodium content, the control solution (Ctrl) was matched for sodium concentration. Colors indicate luminal diameter (red, constricted; blue, relaxed). **(B)** Production of phenethylamine (PEA) by *R. gnavus* in minimal medium supplemented with phenylalanine and tyrosine. Values are normalized to optical density to account for differences in bacterial growth. **(C)** Quantification of contractile activity normalized to baseline. Contractile strength is expressed as the difference between contraction and relaxation phases. The right panel shows the dose-response relationship for PEA-induced tonic contraction, yielding an $EC_{50}$ of 50-75 µM. These data indicate a physiologically relevant concentration range consistent with levels observed in donor-derived fermentations.
